# Structure-Aware Dual-Target Drug Design through Collaborative Learning of Pharmacophore Combination and Molecular Simulation

**DOI:** 10.1101/2023.12.10.571029

**Authors:** Sheng Chen, Junjie Xie, Renlong Ye, David Daqiang Xu, Yuedong Yang

**Author notes:** S.C, Y.Y, and D.D.X designed research; S.C, J.X, and R.Y performed research; S.C, R.Y contributed new reagents/analytic tools; S.C, J.X, and R.Y analyzed data; S.C, Y.Y, D.D.X, J.X, and R.Y wrote the paper.

## Abstract

Dual-target drug design has gained significant attention in the treatment of complex diseases, such as cancers and autoimmune disorders. A widely employed design strategy is combining pharmacophores to incorporate the knowledge of structure-activity relationships of both targets. Unfortunately, it often struggles with long and expensive trial and error, because protein pockets of two targets impose complex structural constraints on the pharmacophore combination. In this study, we propose AIxFuse, a structure-aware dual-target drug design method that learns pharmacophore fusion patterns to satisfy the dual-target structural constraints simulated by molecular docking. We utilize two self-play reinforcement learning (RL) agents to learn pharmacophore selection and fusion by comprehensive feedback including dual-target molecular docking scores. Collaboratively, the molecular docking scores are learned by active learning (AL). Through collaborative RL and AL, AIxFuse learns to generate molecules with multiple desired properties. AIxFuse is shown to outperform state-of-the-art methods in generating dual-target drugs against glycogen synthase kinase-3 beta (GSK3*β*) and c-Jun N-terminal kinase 3 (JNK3). When applied to another task against retinoic acid receptor-related orphan receptor *γ*-t (ROR*γ*t) and dihydroorotate dehydrogenase (DHODH), AIxFuse exhibits consistent performance while compared methods suffer performance drops, leading to a 5 times outperformance in success rate. Docking studies demonstrate that AIxFuse can generate molecules concurrently satisfying the binding mode required by both targets. Further free energy perturbation calculation indicates that the generated candidates have promising binding free energies against both targets.

**Significance Statement:** Complex diseases like cancers and autoimmune disorders are mostly caused by multiple genes. Designing dual-target drugs against two target proteins simultaneously can achieve synergistic effects and alleviate drug resistance. In this study, we present AIxFuse, which to our knowledge is the first structure-aware dual-target drug design method that learns pharmacophore fusion patterns to satisfy the dual-target structural constraints simulated by molecular docking. AIxFuse exhibits superior performance to previous state-of-the-art methods on comprehensive benchmarks. By generating diverse drug candidates with promising dual-target binding free energies and other desired properties, AIxFuse holds promising prospects for accelerating the development of novel dual-target drugs for long-term therapeutic of complex diseases.

---

The paradigm of targeted-drug design has been dominated by the “one target, one drug”(1). However, single-target drugs are prone to drug resistance. For complex diseases like cancers and autoimmune disorders, current single-target therapy struggles to achieve long-term therapeutic effects(2, 3). To engage different therapeutic targets simultaneously, combination therapy has been widely employed to produce additive or synergistic effects by using multiple drugs that act on different targets(4–6). Nevertheless, using multiple drugs involves drug-drug interactions, dose-limiting toxicities, unpredictable pharmacokinetics, adverse off-target effects, and poor patient compliance(7). A potential way to overcome these limits is to design dual/multi-target drug (8–10). In recent years, there has been an increasing trend towards dual/multi-target drugs in FDA-approved medications, particularly in the treatment of malignant tumors, nervous system diseases, as well as digestive and metabolic systems diseases(11, 12).

Towards rational design of dual-target drugs, the pharmacophore combination strategy has been widely employed(10, 13–17). It includes simple “linking” of two distinct pharmacophores with a linker, “merging” of pharmacophores with shared fragments, and “fusing” of pharmacophores. The “linking” approach often results in higher molecular weight and is prone to suffering from pharmacokinetic issues(10, 13). The “merging” approach is limited to targets that share some binding modes (14, 15). The “fusing” approach allows tight joining of active fragments and offers the opportunity to achieve desired physicochemical properties in the resulting molecules(16, 17). Designing dual-target drugs by pharmacophore combination can effectively integrate the knowledge of structure-activity relationships (SAR) on both targets (2). Unfortunately, it often struggles with long and expensive trial and error because the protein pockets of two targets impose complex structural constraints on the pharmacophore combination patterns. Therefore, it is valuable to develop dual-target drug design methods that improve the efficiency and success rate.

Computational drug design methods have been developed over decades to generate molecules with desired properties(18, 19). Traditional molecular generation primarily navigated the chemical-space exploration through population-based stochastic optimization procedures, such as evolutionary algorithms (EA) (20) or swarm intelligence (21). In recent years, deep learning algorithms have been widely employed in molecular generation (22, 23). Early studies such as ChemVAE (24), CVAE (25) and Heteroencoder (26) generated SMILES strings as molecular representation. On the other hand, Liu et al. (27) and You et al. (28) made exploratory attempts to generate molecules represented by the graph, whose node-edge structure naturally matches the atom-bond structure of molecules. Unfortunately, most of them focused on none/single-objective generation and thus are not suitable for dual-target drug design. There are only a handful of multi-objective molecular generation algorithms (29–33). Li et al.(33) trained machine learning (ML) models as approximate empirical measurements of activity against glycogen synthase kinase-3 beta (GSK3*β*) and c-Jun N-terminal kinase 3 (JNK3). Following their work, related methods (e.g. RationaleRL(31), MARS(29)) employed the same activity predictors in their methods and assessments. However, it brought up two issues. Firstly, the limited publically available experimental activity data could not guarantee the generalizability of the ML model. Secondly, The ML model based on molecular fingerprint representation is not structurally interpretable. Compared with the ML-based activity predictor, molecular-simulation-based computational tools such as molecular docking or free energy perturbation (FEP) are more generalizable and structurally interpretable. But they are less computationally efficient. Recent efforts have been made on molecular docking (34, 35) and FEP (36, 37) by active learning (AL) to improve the efficiency of virtual screening. Therefore, integrating multi-objective molecular generation with active learning on molecular-simulation-based activity estimation is a promising way to improve generalizability and structural interpretability.

However, combining molecule generation with AL does indeed present a unique challenge. In the context of virtual screening, the target domain for AL is well-defined, given that the compound library is known and readily available. In contrast, when it comes to molecule generation, the target domain for AL remains elusive because the molecules within this domain haven’t been generated. This dilemma is akin to training an artificial intelligence algorithm for the GO game in the absence of any historical chess records. AlphaZero (38) has masterfully navigated this very conundrum by self-play with reinforcement learning (RL) and Monte Carlo tree search (MCTS). Inspiring, searching pharmacophore combination patterns to satisfy the dual-target structural constraints can be also modeled as two self-play MCTS agents against two targets. Through self-play pharmacophore combination, target domain molecules can be generated for AL training, and the trained models can subsequently feedback to navigate the molecular generation.

In this study, we propose AIxFuse, which to our knowledge is the first structure-aware dual-target drug design method that learns pharmacophore fusion patterns to satisfy the dual-target structural constraints simulated by molecular docking. In AIxFuse, pharmacophores are extracted automatically through protein-ligand interaction analysis on active compounds and target proteins. Molecular sub-structure trees are built for both targets to represent the pharmacophore-fusion chemical space. Exploration in this chemical space towards multiple desired properties (i.e. binding affinity, drug-likeness, and synthetic accessibility) is navigated by an Actor-Critic-like(39) RL framework. The RL framework consists of two self-play MCTS Actors for molecule generation and a dual-target docking score Critic trained by AL. After iterations of collaborative RL and AL, AIxFuse learns to generate molecules with multiple desired properties. AIxFuse was first evaluated on designing dual-inhibitor for GSK3*β* and JNK3, where it showed outperformance (32.3% relative improvement in success rate) compared to other state-of-the-art (SOTA) methods. We also applied AIxFuse to another task that aims to design dual inhibitors against retinoic acid receptor-related orphan receptor *γ*-t (ROR*γ*t) and dihydroorotate dehydrogenase (DHODH) (3), where AIxFuse achieves a success rate of 23.96%, over 5 times higher than the best of other methods. Docking studies demonstrated that AIxFuse can generate molecules that concurrently satisfy the binding mode required by both targets. Furthermore, as revealed by FEP, a generated candidate exhibits better/comparable binding free energy with known inhibitors on both targets, demonstrating its dual-target activity potential. This holds promising prospects for AIxFuse to accelerate dual-target drug design in real-world drug development scenarios.

## Results

Fig. 1 displays the overall pipeline of AIxFuse. For both targets, the targeting protein structures and known active compounds are first collected and their protein-ligand complex structures are simulated by molecular docking. From the complex structures, protein-ligand interactions (PLIs) are extracted through the PLIP(40, 41) program. The extracted PLIs are then combined to define the pharmacophores (Fig. 1B) that are ranked by interaction scores, frequency scores, etc. The top-scored pharmacophores split active compounds into core and side chain fragments. These fragments can be reorganized by two searching trees (Fig. 1C), whose structures are detailed in the Materials and Methods section. The leaf nodes of two trees will be fused to generate the pharmacophore-fused molecules. An Actor-Critic-like reinforcement learning (RL) framework is employed to learn the optimal pharmacophore fusion patterns. It consists of two self-play Monte Carlo Tree Searching (MCTS) Actors and a dual-target docking score Critic. More specifically, MCTS Actors aim to maximize the upper confidence bounds (UCB) of the reward for the generated molecules. The reward function encompasses various properties, including drug-likeness, synthetic accessibility, and dual-target docking scores. As a dual-target docking score Critic, a multi-task AttentiveFP(42) model is iteratively trained by the active learning (AL). After several iterations of collaborative RL and AL, the final models will be used to generate molecules.

**Fig. 1.**
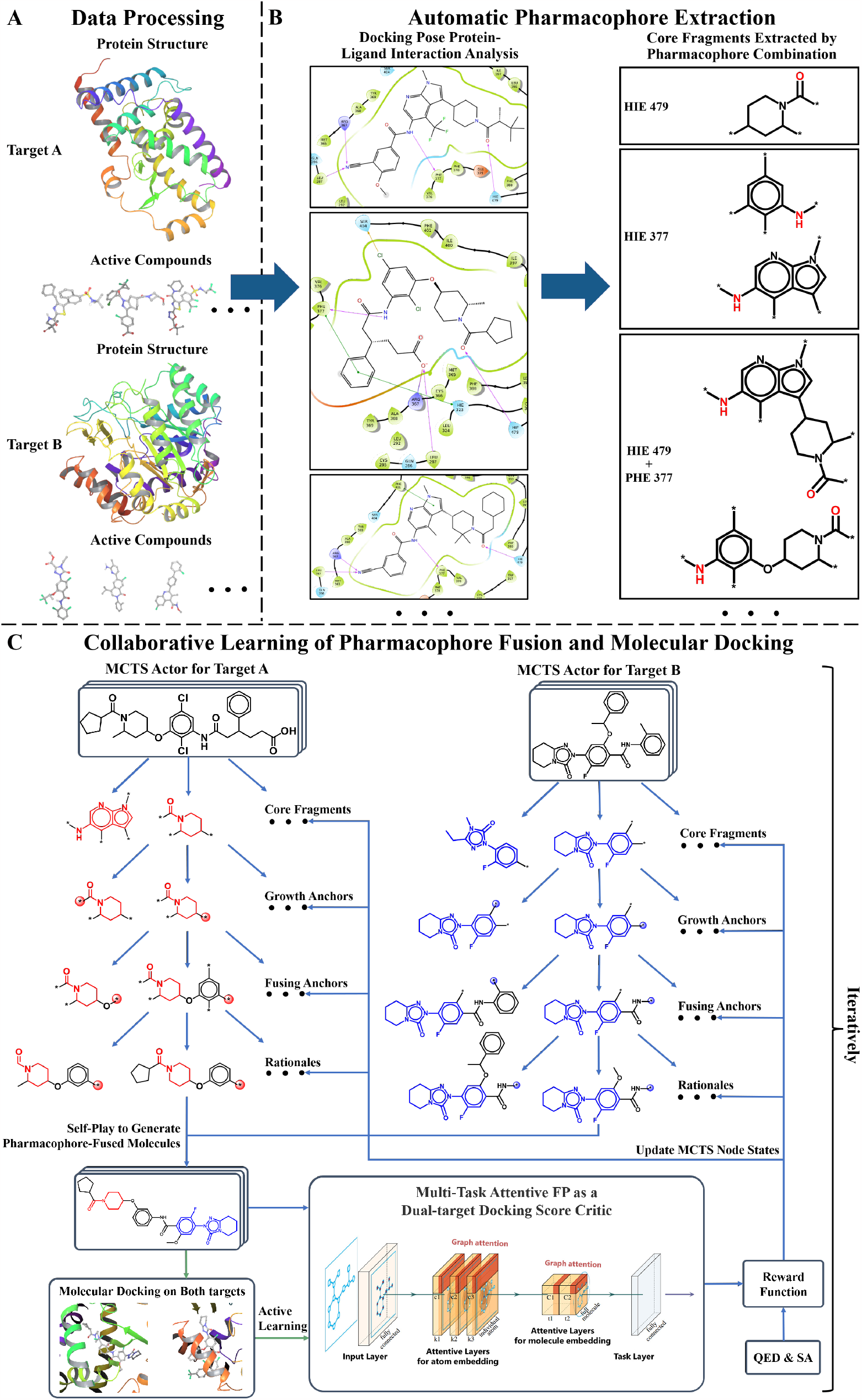
The AIxFuse pipeline encompasses three core stages: (A) Data Processing, entailing the preprocessing of protein structures and active compounds for both targets; (B) Automatic Pharmacophore Extraction, accomplished through automatic scrutiny of protein-ligand interactions within docking poses of active compounds to isolate core fragments that preserve key pharmacophores; and (C) Collaborative Learning of Pharmacophore Fusion and Molecular Docking. The Actor-Critic-like reinforcement learning on pharmacophore fusion and the active learning on molecular docking are run collaboratively and iteratively. It employs two self-play MCTS Actors for molecule generation and a dual-target docking score Critic to navigate generation. The Critic is a multi-task AttentiveFP (schematic diagram depicted by Xiong et al.’s (42)) and is trained by active learning. The green arrow lines represent the training procedure.

### AIxFuse Outperformed Previous SOTA Methods on GSK3*β* | JNK3 Dual-inhibitor Design

AIxFuse was compared to state-of-the-art (SOTA) methods: RationaleRL(31), MARS(29), and REINVENT2.0(32). We ran each method to generate 10,000 molecules for designing dual inhibitors against glycogen synthase kinase-3 beta (GSK3*β*) and c-Jun N-terminal kinase 3 (JNK3), a classical benchmark for dual-target drug design(29, 31, 33).

All methods were first assessed in terms of three general metrics for molecular generation: validity, uniqueness, and diversity. As shown in Table 1, all methods achieved a validity rate exceeding 98%, signifying their ability to generate molecules that conform to essential chemical constraints. Notably, significant variations were observed in the uniqueness metric. AIxFuse demonstrated the highest uniqueness at **89.7%**, surpassing previous other methods, including REINVENT2.0 (82.6%), RationaleRL (53.3%), and MARS (24.1%). Although fragment-based methods often struggle with diversity, AIxFuse obtained a score of 0.719, outperforming RationaleRL (0.656) and MARS (0.597), and matching REINVENT2.0 (0.722). This unexpected success may be attributed to the upper confidence bounds (UCB) of reward expectations in MCTS, which aims to strike a balance between sampling frequency and reward expectations.

**Table 1.**
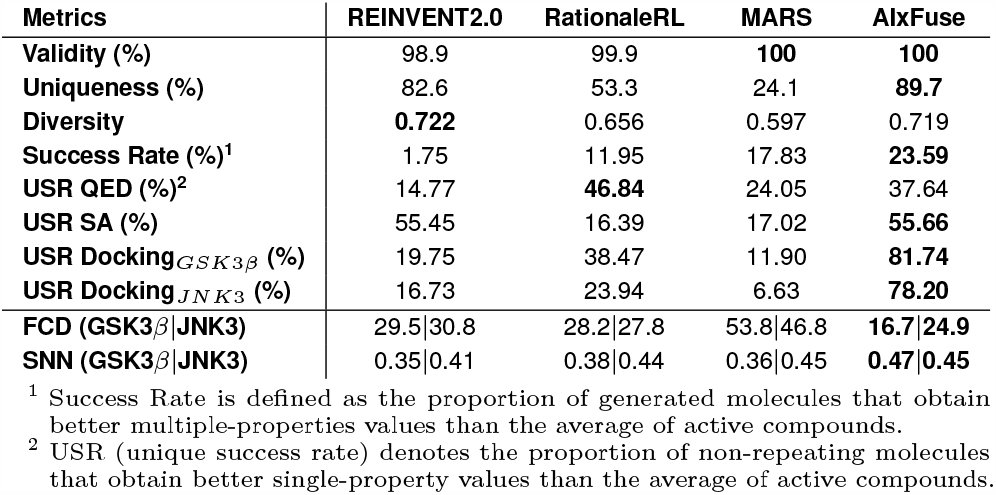
Performance of AIxFuse and compared methods on GSK3*β* | JNK3 dual-target drug design benchmark.

We then checked the ability of each method to generate molecules with multiple desired properties. The success rate is defined as the proportion of generated molecules that obtain better dual-target docking scores, Quantitative Estimate of Drug-likeness (QED), and synthetic accessibility (SA) than the mean values of known active compounds. As shown in Table 1, AIxFuse achieved the highest success rate at **23.59%**, outperforming the second-place method MARS (17.83%) with a relative improvement of **32.3%**. It demonstrated AIxFuse’s ability to generate molecules that simultaneously satisfy multiple property constraints. To focus on each single property, the unique success rate (USR) was calculated as the proportion of non-repeating molecules satisfying an individual property constraint. By replacing RL and AL with random sampling on both trees, we constructed AIxFuse(w/o RLAL) as a baseline model. As shown in Fig. 2, baseline model AIxFuse(w/o RLAL) exhibited promising USR in docking scores, but its USRs in SA and QED were not optimal. By incorporating RL and AL, AIxFuse achieved the best USR in SA (55.66%) and the second-highest USR in QED (37.64%), demonstrating significant improvements to AIxFuse(w/o RLAL). Furthermore, AIxFuse achieved the highest USR on the docking scores of GSK3*β* (81.74%) and JNK3 (78.20%), surpassing other methods by a significant margin.

**Fig. 2.**
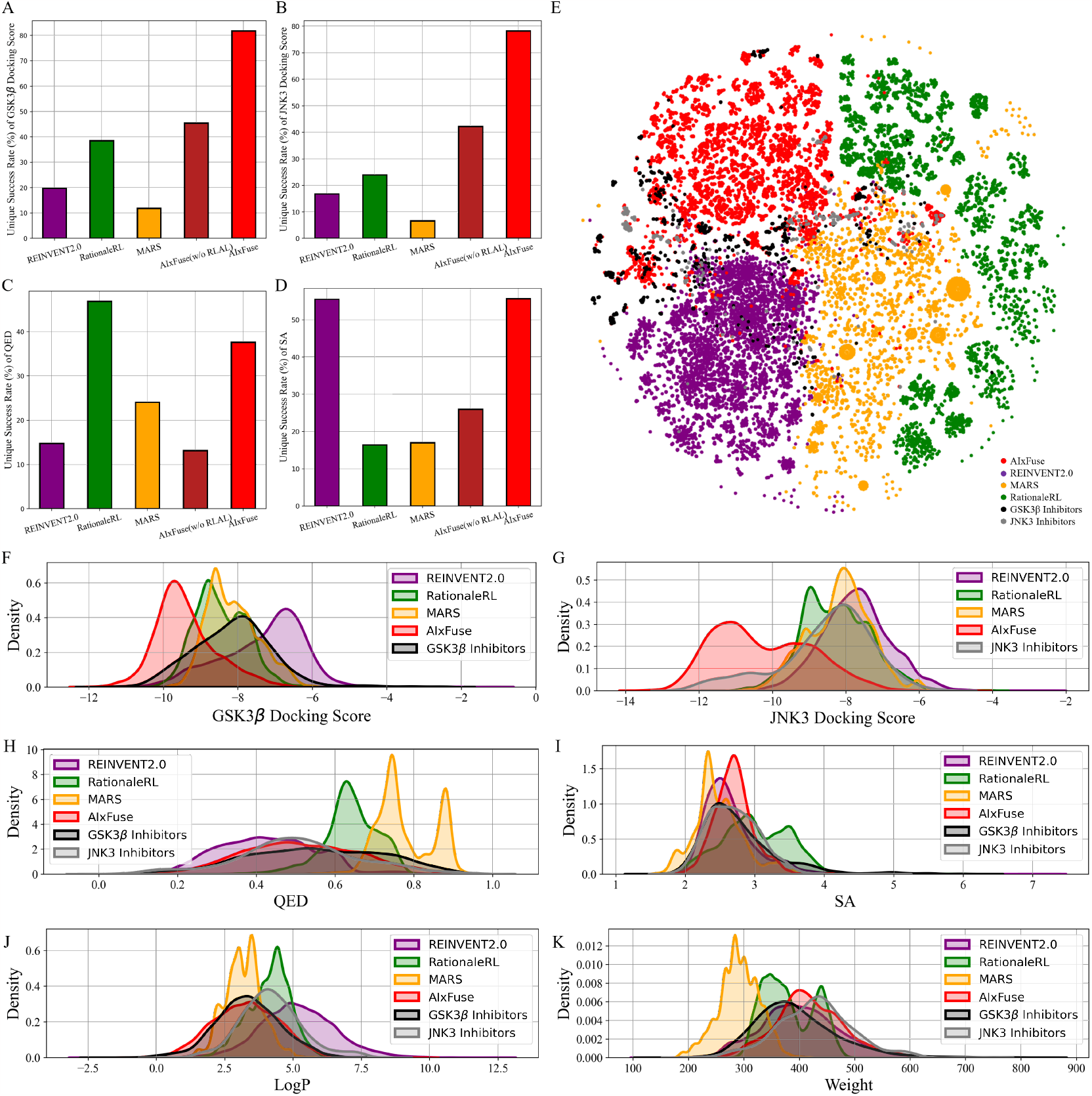
Statistics and visualization of molecules generated for GSK3*β* JNK3 dual-inhibitor design: The unique success rate of (A) GSK3*β* docking score, (B) JNK3 docking score, (C) QED, and (D) SA; (E) The t-SNE visualization of Morgan fingerprints of known inhibitors and generated molecules; The property distributions of (F) GSK3*β* docking score, (G) JNK3 docking score, (H) QED, (I) SA, (J) LogP, and (K) molecular weight.

We were interested in studying the similarity between generated molecules and active compounds. As shown in Table 1, AIxFuse obtained the lowest Fréchet ChemNet Distance (FCD) and the highest Similarity to Nearest Neighbor (SNN) against active compounds of both targets consistently. This is as expected since AIxFuse preserved key pharmacophores of active compounds to incorporate SARs of both targets. Towards an intuitive understanding of similarity, Fig. 2E visualized the chemical spaces of generated molecules and active compounds. The chemical space of the GSK3*β* inhibitors was close to that of the JNK3 inhibitors. Notably, AIxFuse explored the gaps between main clusters of known inhibitors, while other methods didn’t. Molecules generated within the chemical space gaps between inhibitors of two targets are likely to exhibit high similarity to both ends. Looking deeper into the molecular property similarity between generated molecules and active compounds, we plotted the property distributions in Fig. 2F-K. It can be found that the QED, SA, LogP, and Weight distribution of AIxFuse-generated molecules closely aligned with those of GSK3*β* inhibitors and JNK3 inhibitors. However, for both GSK3*β* and JNK3 docking scores, AIxFuse-generated molecules were distributed to the lower end of the spectrum than corresponding known inhibitors. It demonstrated that AIxFuse-generated molecules obtained better estimated binding affinities against both targets while keeping other properties similar to known inhibitors.

What contributed to the outperformance of AIxFuse? We first explored the contribution of running multiple iterations of AL. As shown in Fig. 3A-B, as AL iterated, the overall MSE decreased while R^2^ increased. This trend demonstrated that multiple iterations of AL contribute to the accuracy of dual-target docking score Critic. We then presented the accuracy of the final dual-target docking score Critic in Fig. 3C-D. The final Pearson correlation coefficients (0.691 and 0.704 for GSK3*β* and JNK3, respectively) indicated the strong positive correlation between the predicted docking scores and the ground truth. This affirmed the models’ robust ranking ability, which is an essential factor for molecular generation to improve molecular properties. We then investigated the contribution of RL optimization. As is shown in Fig. 3E-H, the docking score distribution of AIxFuse(w/o RLAL) closely resembles that of known active molecules. This suggests that our automatically extracted pharmacophores can effectively preserve essential SARs of both targets. However, AIxFuse(w/o RLAL)’s QED distribution and SA distribution are not ideal. After iterations of collaborative RL and AL, AIxFuse achieves improvements in all four properties. These phenomenons underscore the synergistic effect of collaborative RL and AL: AL trained dual-target docking score Critic for RL optimization, while RL provided molecules with higher quality for AL training.

**Fig. 3.**
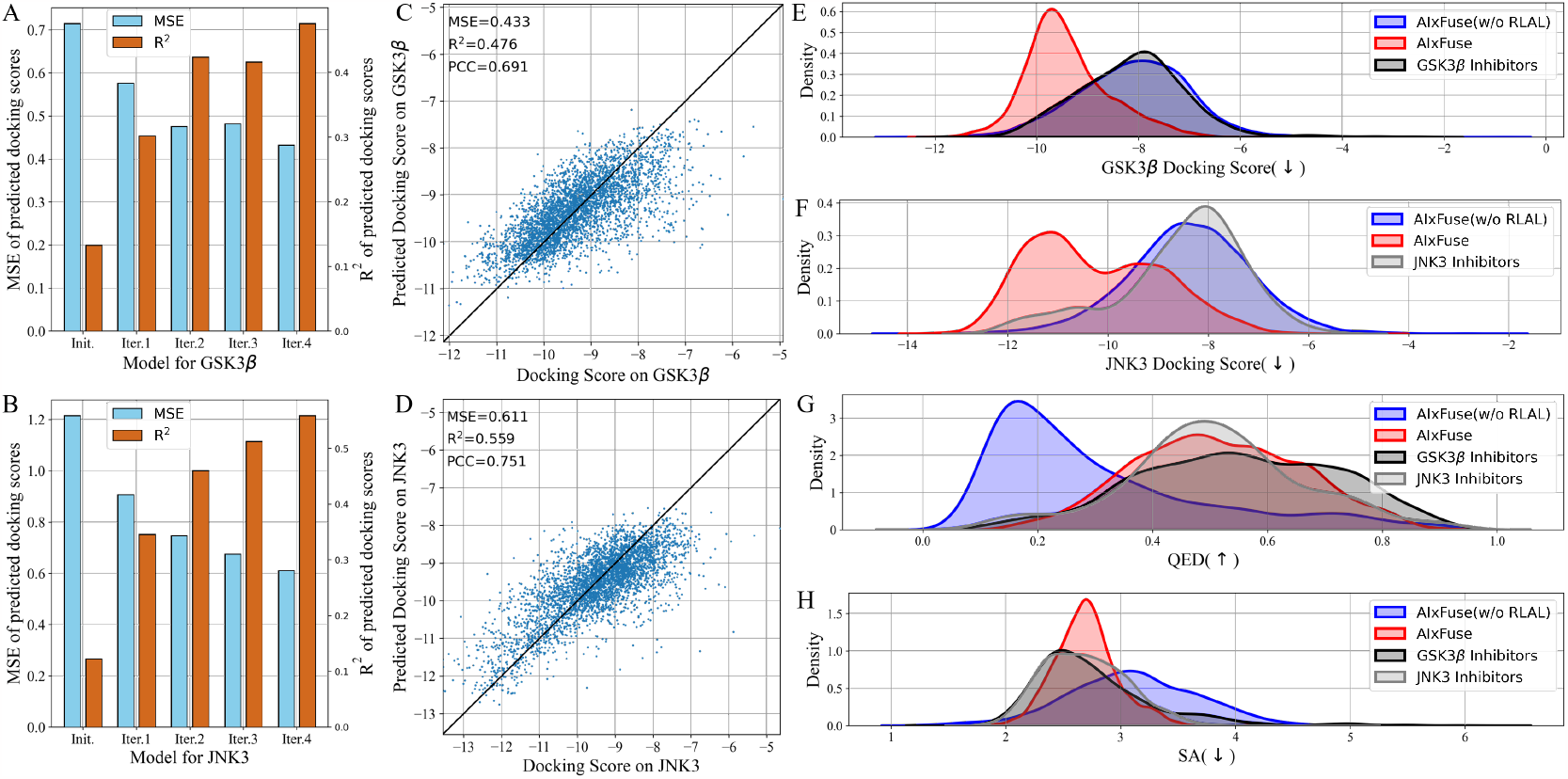
The ablation study of multiple iterations of AL: The MSE and R^2^ of docking scores on (A) GSK3*β* and (B) JNK3 predicted by models at various iteration steps; The scatter plot illustrating the correlation between (C) GSK3*β* and (D) JNK3 docking scores and predicted docking scores at final iteration; The ablation study of collaborative RL and AL: The property distribution of (E) GSK3*β* docking scores, (F) JNK3 docking scores, (G) QED, and (H) SA of molecules generated by AIxFuse(w/o RLAL), AIxFuse, and active compounds of GSK3*β* and JNK3.

### AIxFuse Achieved Consistent Outperformance on ROR*γ*t|DHODH Dual-inhibitor Design

We then applied AIxFuse to another dual-target design task, where there is a precedent for rational design. Chen et al. previously designed a potential dual-target inhibitor **(R)-14d**(3), achieving IC50 values of 0.110*μ*M for ROR*γ*t (retinoic acid receptor-related orphan receptor *γ*-t) and 0.297*μ*M for DHODH (dihydroorotate dehydrogenase). Following the evaluation setting of GSK3*β*| JNK3 benchmark task, we ran AIxFuse, RationaleRL, MARS, and REINVENT2.0 to generate 10,000 molecules for ROR*γ*t | DHODH dual-inhibitor design.

Compared with other methods, as shown in Table 2, AIxFuse achieved the highest validity, uniqueness, success rate, USR QED, USR SA, USR Docking_*RORγt*_, USR Docking_*DHODH*_, USR 3D SNN_*RORγt*_, and USR 3D SNN_*DHODH*_ . The diversity achieved by AIxFuse is lower than REINVENT2.0, comparable with RationaleRL, and far higher than MARS. When compared across benchmark tasks, AIxFuse exhibited consistent results, whereas other methods displayed fluctuations. Fig. 4A highlights that AIxFuse achieved a success rate of 23.96%, consistent with its performance in the GSK3*β* | JNK3 task (23.59%, Fig. 4B). Conversely, MARS failed to generate any successful molecules against ROR*γ*t | DHODH, resulting in a success rate drop (17.83% → 0%). A similar decrease in success rate occurred in RationaleRL (11.95% → 1.16%). Both MARS and RationaleRL rely on machine learning (ML) activity predictors. However, due to the limited availability of public non-active data for DHODH, training ML predictor posed challenges (detailed in *Supplementary Text 1*). In contrast, AIxFuse replaced ML-based activity predictors with molecular docking, which should contribute to its consistent performance.

**Table 2.**
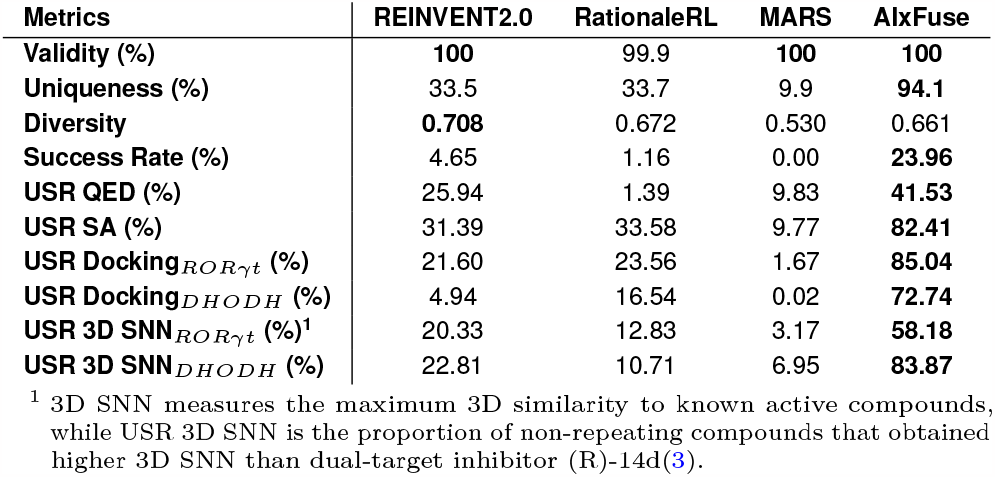
Performance of AIxFuse and compared methods on ROR*γ*t | DHODH dual-target drug design benchmark.

**Fig. 4.**
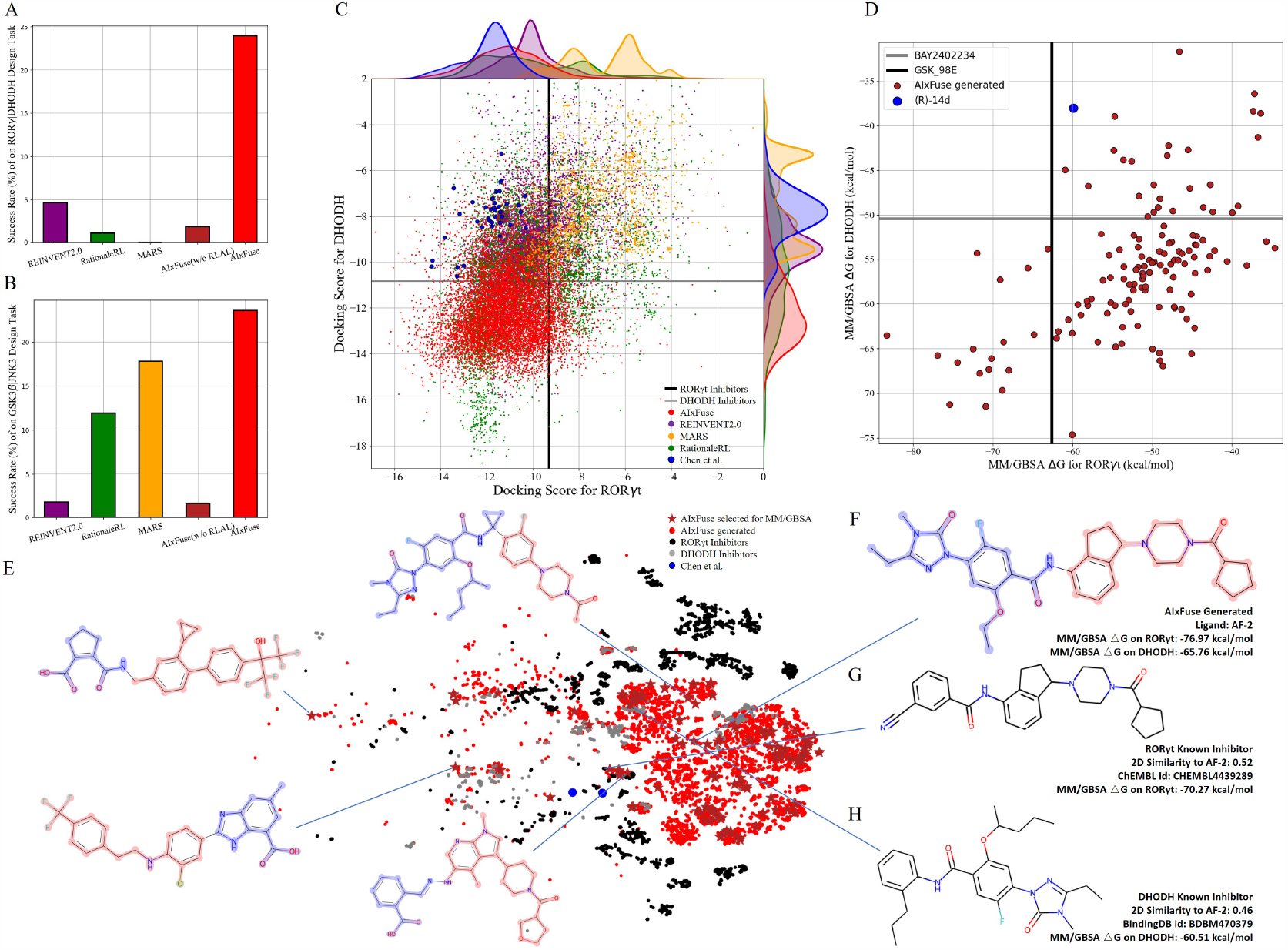
The performance of AIxFuse on ROR*γ*t DHODH dual-inhibitor design: The success rates of AIxFuse and compared methods on (A) ROR*γ*t DHODH dual-inhibitor design with (B) GSK3*β* JNK3 task as a reference; The scatter plot of dual-target docking scores (C) obtained by AIxFuse (red), REINVENT2.0 (purple), MARS (yellow), and RationaleRL (green) with compounds designed by Chen et al. as references (blue); (D) The dual-target MM/GBSA binding free energies of 134 AIxFuse-generated molecules (brick red) with dual-target inhibitor (R)-14d (blue) as a reference; The t-SNE visualization (E) of AIxFuse-generated molecules that is selected for MM/GBSA calculation with molecular structure of some examples; (F) The molecular structure and MM/GBSA binding free energies of AIxFuse-generated drug candidate AF-2 with its most similar ROR*γ*t inhibitor (G) CHEMBL4438289 and its most similar DHODH inhibitor (H) BDBM470379.

We then checked the dual-target activity potential of AIxFuse-generated molecules. As shown in Fig. 4C, AIxFuse’s docking score distributions on both targets consistently trend towards the lower (better) end of the spectrum. Most AIxFuse-generated molecules exhibited lower ROR*γ*t and DHODH docking scores than the average score of known ROR*γ*t inhibitors and DHODH inhibitors respectively, indicating promising dual-target activity potential. When referring to compounds designed by Chen et al., AIxFuse also successfully generated numerous molecules with comparable ROR*γ*t docking scores and lower DHODH docking scores. Molecular Mechanics / Generalized Born Surface Area calculation (MM/GBSA) provides a more accurate estimation of protein-ligand binding free energies(43). 134 AIxFusegenerated molecules were selected (detailed in the Materials and Methods section) for MM/GBSA calculation. Fig. 4D illustrates their computation results. When referring to the GSK 98E (active ligand in the co-crystal structure of ROR*γ*t, PDB id: 5NTP(44)), 17 AIxFuse-generated molecules obtained lower binding free energies. All these 17 candidates achieved lower binding free energies than BAY2402234 (active ligand in the co-crystal structure of DHODH, PDB id: 6QU7(45)). We then performed t-SNE visualization of the chemical space of these selected molecules in Fig. 4E and displayed structures of 5 representative molecules, which showed good structural diversity. Looking into a detailed case, AF-2 (structure shown in Fig. 4F) exhibited dual-target activity potential with MM/GBSA binding free energies of -76.97 *kcal/mol* on ROR*γ*t and -65.76 *kcal/mol* on DHODH. We also displayed structures of AF-2’s most similar ROR*γ*t inhibitor (ChEMBL id: CHEMBL4438289) and most similar DHODH inhibitor (BindingDB id: BDBM470379) in Fig. 4G and Fig. 4H, respectively. Notably, the left half structure (LHS) of CHEMBL4438289 showed a similar pattern to the LHS of BDBM470379. AF-2 successfully merged this pattern by fusing the ROR*γ*t rationale (red in Fig. 4F) and the DHODH rationale (blue in Fig. 4F). This should contribute to its lower MM/GBSA binding free energies on both targets.

### Binding Mode and Affinity Study on the AIxFuse-Generated Dual-Target Drug Candidate against ROR*γ*t | DHODH

Absolute protein-ligand binding free-energy calculation by free energy perturbation (ABFEP)(46) is one of the most reliable in-silico approaches. ABFEP calculation has achieved a remarkable root mean square error of 1.1 *kcal/mol* after zero-point shifting to match the experimental values in a previous study (47). Therefore we employed ABFEP calculation as the last process of molecular screening. As shown in Fig. 5A, we selected 17 molecules with balanced dual-target MM/GBSA binding free energies for ABFEP calculations. Among them, we found AF-5, AF-20, AF-2, and AF-16 with binding free energies comparable to those of the active molecules for both targets (detailed in *Supplementary Table S6*). Finally, AF5 was chosen as the top dual-target drug candidate as it exhibited the most promising dual-target activity potential.

**Fig. 5.**
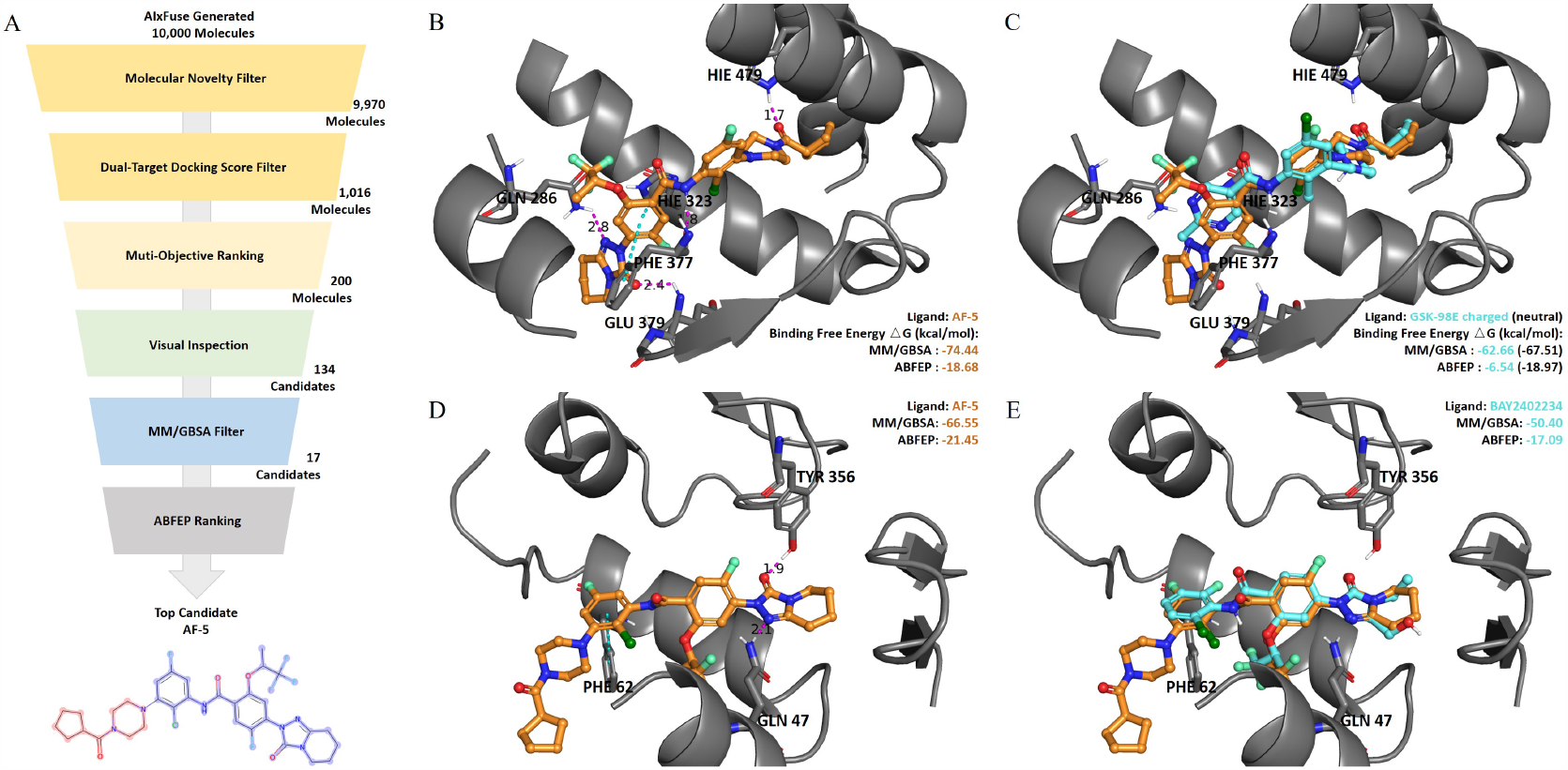
(A) The virtual screening process on AIxFuse-generated molecules. (B) The binding mode of AIxFuse generated compound AF-5 (orange stick) with ROR*γ*t-LBD revealed by docking study. (C) Overlay of AF-5(orange stick) with ROR*γ*t-LBD co-crystal ligand GSK-98E (cyan stick). (D) Binding mode of AF-5(orange stick) with DHODH. (E) Overlay of AF-5(orange stick) with DHODH co-crystal ligand BAY2402234 (cyan stick). The pink dotted lines represent hydrogen bonds, and the cyan dotted lines represent *π*-*π* stacking.

We first studied the binding mode of AF-5 on both targets by molecular docking. As shown in Fig. 5B, AF-5 established hydrogen bonds and *π*-*π* stacking interactions with key residues, including HIE479, PHE377, GLU379, HIE323, and GLN 286. we then docked AF-5 to another target DHODH. As shown in Fig. 5D, AF-5 exhibited the hydrogen bonds and *π*-*π* stacking between AF-5 and key residues GLN47, TYR356, and PHE62. AF-5’s binding modes on both targets matched previous studies on corresponding known inhibitors (3, 44, 45). Fig. 5C and Fig. 5E illustrated that the docking pose of AF-5 (colored in orange) on both targets closely resembled those of two known inhibitors (ROR*γ*t inhibitor GSK-98E in the co-crystal structure 5NTP, and DHODH inhibitor BAY2402234 in 6QU7, both colored in cyan).

We then analyzed the dual-target activity potential of AF-5 by ABFEP. AF-5 was compared with ROR*γ*t inhibitor GSK-98E and DHODH inhibitor BAY2402234 on their binding free energies against the corresponding target. According to its optimal protonation state at pH 7.0 *±* 2.0, GSK-98E had a +1 net charge, while AF-5 was electrically neutral. The difference in net charge would cause systematic errors. Therefore, we also calculated the binding free energy of the GSK-98E in the electrically neutral state. As shown in Fig. 5C, AF-5’s ROR*γ*t binding free energy (-18.68 *kcal/mol*) was lower than that of charged GSK-98E (-6.54 *kcal/mol*) and comparable to that of neutral GSK-98E (-18.97 *kcal/mol*). Against another target DHODH, the AF-5’s ABFEP binding free energy (-21.45 *kcal/mol*) was much lower than that of BAY2402234 (-17.09 *kcal/mol*). We also supplemented the ABFEP calculation results of several other active molecules as additional references. As shown in Supplementary Table S6, AF-5’s ROR*γ*t binding free energy was comparable to those of known ROR*γ*t inhibitors, while its DHODH binding free energy was lower than those of known DHODH inhibitors. Insilico, it demonstrated the potential of AF-5 as a dual-target inhibitor against ROR*γ*t and DHODH.

## Discussion

Dual-target drug design is attractive but still challenging. Traditional pharmacophore-combination-based manual design often struggles with long and expensive trial and error. This is because the protein pockets of two targets impose complex structural constraints on the pharmacophore combination. Therefore, it is important to develop computational methods that learn pharmacophore fusion patterns to satisfy the dualtarget structural constraints. AIxFuse, as an innovative computational dual-target drug design method, has made solid progress on the challenging problem of improving the generalizability and structural interpretability of dual-target molecular generation. By utilizing molecular docking rather than ML-based activity prediction, AIxFuse exhibited enhanced generalizability (independent of the size of the training dataset). AIxFuse was also structurally interpretable, as its pharmacophores were extracted based on protein-ligand interaction and it optimized the target-specific binding affinities by molecular docking. AIxFuse turns pharmacophore fusion and molecular docking learnable through iterations of collaborative reinforcement learning and active learning. To our knowledge, AIxFuse is the first structure-aware dual-target drug design method that learns pharmacophore fusion patterns to satisfy the dual-target structural constraints simulated by molecular docking.

AIxFuse was benchmarked on two dual-target drug design tasks and compared with previous SOTA methods. In the dual-target drug design task against GSK3*β* and JNK3, AIxFuse exhibited 32.3% relative improvement in success rate compared with the best of other methods. In the ROR*γ*t and DHODH dual-target inhibitor design task with limited activity data, AIxFuse achieves a success rate of 23.96% while compared methods suffered performance drops, leading to a 5 times outperformance in success rate. Further in-silico binding free-energy calculation results highlighted the dual-target activity potential of AIxFuse-generated drug candidates.

As a structure-aware dual-target drug design method, AIxFuse can generate potential dual-target hit compounds for subsequent lead optimization. For example, our latest structure-based scaffold decoration method DiffDec(48) can be used for lead optimization on AIxFuse-generated molecules. In the future, active learning on FEP(36, 37) can be also introduced into AIxFuse to enhance the accuracy of binding affinity estimation. As the demand for dual-target drugs becomes significant in the treatment of complex diseases, AIxFuse can provide rapid and effective starting points for various dual-target design tasks.

## Materials and Methods

### Benchmark Curation

To assess the efficacy of our method, we established two dual-target inhibitor design benchmark tasks. The first benchmark involved the development of dual-target inhibitors against Glycogen synthase kinase-3 beta (GSK3*β*) and c-Jun N-terminal kinase 3 (JNK3), denominated as the GSK3*β* | JNK3 benchmark. Notably, GSK3*β* and JNK3 have been associated with the pathogenesis of Alzheimer’s disease (AD) and other disorders (49, 50). The concept of designing dual inhibitors concurrently against GSK3*β* and JNK3 holds the potential to introduce a novel multi-target therapy for AD (33).

The second benchmark task aims to design dual-target inhibitors against retinoic acid receptor-related orphan receptor *γ*-t (ROR*γ*t) and dihydroorotate dehydrogenase (DHODH), named as the ROR*γ*t—DHODH benchmark. ROR*γ*t and DHODH have both emerged as attractive targets for the treatment of autoimmune diseases (51, 52). Chen et al. (3) advocated for the development of ROR*γ*t | DHODH dual inhibitor as a promising strategy to enhance therapeutic efficacy, reduce toxicity, and mitigate drug resistance in the context of inflammatory bowel disease (IBD) therapy. They designed a series of compounds and identified one candidate with both in vitro and in vivo activities, along with favorable mouse pharmacokinetic profiles.

### Evaluation Settings

On the above two benchmarks, We compared AIxFuse with the following methods, and their implementation details are available in *Supplementary Text 4* :

- RationaleRL(31): RationaleRL begins by extracting rationales from active compounds via MCTS to retain essential activity-contributing elements. Subsequently, it employs the pharmacophore merge strategy to combine rationales, preserving the maximum common substructure. Finally, a molecular generation model is fine-tuned to produce side chain groups.
- MARS(29): MARS selects the top 1000 fragments from the ChEMBL dataset, ensuring that each fragment contains no more than 10 heavy atoms and appears most frequently. It employs Graph Neural Networks (GNN) and Markov chain Monte Carlo (MCMC) sampling to manipulate these fragments and generate molecules with enhanced properties.
- REINVENT2.0(32): This method generates SMILES representations of molecules using a Recurrent Neural Network (RNN). Reinforcement learning is then applied to optimize the properties of the generated molecules, facilitating the generation of multi-target molecules through the application of multiple rewards.

We employed a diverse set of metrics to assess the performance of each method. Initially, we computed general metrics such as validity, uniqueness, and diversity using Moses(53). Additionally, we scrutinized properties including drug-likeness, and synthesizability of the generated molecules and dual-target activity. Quantitative Estimate of Drug-likeness (QED) and synthetic accessibility (SA) were calculated using RDKit(54). Notably, for dual-target activity estimation, no public ML-based active predictor is available for the ROR*γ*t | DHODH task. Therefore, we opted for the molecular docking score as the approximation of binding affinity in both tasks. As a molecular-simulation-based method, molecular docking offers structural interpretability and is independent of the quantity of available activity data. Following the approach of Chen et al.(3), Glide (55) software was employed to estimate binding modes and affinities. The co-crystal structures of ROR*γ*t LBD (PDB: 5NTP(44)), DHODH (PDB: 6QU7(45)), GSK3*β*, (PDB: 6Y9S(56)) and JNK3 (PDB: 4WHZ(57)) were selected and processed for docking preparation. The generated molecules were designed to concurrently satisfy multiple property constraints, prompting us to calculate the success rate, denoting the percentage of generated molecules meeting multiple property constraints concurrently. Diverging from prior works(29, 31) which set the thresholds of success as fixed values, we adopted the average value of active compounds as the threshold for each property. Those active compounds were collected from ChEMBL and BindingDB (details available in *Supplementary Text 4*). For the GSK3*β* | JNK3 benchmark, the thresholds for QED, SA, and dual-target docking scores were set at 0.538, 2.76, -8.12, and -8.53 according to the average value of active compounds. For the ROR*γ*t | DHODH benchmark, the corresponding thresholds were 0.430, 3.49, -9.31, and -10.8. To mitigate the contribution of repeated molecules, we also computed the “unique success rate” for each property, which accounts for the proportion of non-repeating compounds that satisfied the constraint. Fréchet ChemNet Distance (FCD)(58) and Similarity to Nearest Neighbor (SNN) scores were calculated by Moses(53) to assess the similarity between generated compounds and known inhibitors. Since designing molecules with higher 3D similarity to lead compounds is desirable in drug design(59), for the ROR*γ*t | DHODH benchmark, we additionally calculated the 3D SNN of the generated molecules by ShapeTanimotoDist in RDKit(54), measuring the maximum 3D similarity to known active compounds.

### Extracting Core Fragments with Key Pharmacophores

The pharmacophore fusing strategy emphasizes the preservation of crucial pharmacophores across diverse targets(2). Here the pharmacophore can be defined as the combination of protein-ligand interactions (PLI). We devised a dedicated module for the extraction of core fragments that retain the essential pharmacophores present in active compounds. This module was designed to facilitate both automatic extraction and guided selection, where expert-recommended key residues can influence the process. In this study, with reference to previous studies(3, 44, 45), we selected specific key residues for ROR*γ*t and DHODH. For ROR*γ*t, HIE479 was set as the key residue. For DHODH, GLN47 and TYR356 were identified as key residues. Regarding GSK3*β* | JNK3, we referred to the binding mode analysis in the co-crystal structures(56, 57), which emphasized the contribution of PLIs associated with VAL135 in GSK3*β*(56) and MET149 in JNK3(57).

To analyze PLIs, we employed the PLIP tool (40, 41) on the molecular docking complex structures due to the unavailability of co-crystal structures for most active compounds. For each pharmacophore, we systematically extracted its minimal connected substructure, termed core fragments, as illustrated in Fig. 1B. Our interest lies in core fragments that possess the following characteristics: 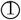 interaction or contact with key residues, 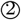 prevalence across multiple active compounds, and 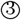 appropriate size. Consequently, for each core fragment, we aggregated its score across all active compounds that share this core. Details of the scoring function are available in *Supplementary Text 2*. Ultimately, the top-scoring core fragments were chosen to construct trees of two targets. The overall pharmacophore extraction and selection process is visualized in Fig. 1B and detailed in *Supplementary Algorithm 1*.

### Tree Structure for Dual-Target Pharmacophore Fusion

Drawing inspiration from the Monte Carlo Tree Search (MCTS) employed in AlphaZero, we were motivated to explore the utilization of tree structures as a representation of the pharmacophore-fusion chemical space. While previous studies have made forays into using tree structures as representations of molecules(60, 61), our approach stands apart in that it introduced a well-defined starting point. This starting point, denoted as **Core Fragments** in Fig. 1C, was derived from the pharmacophore extraction module. Consequently, the initial tier of our tree structure was dedicated to the selection of Cores, which exerted control over the preservation of specific key pharmacophores.

Once two Cores have been selected from two trees, the challenge at hand transforms into how to modify and fuse them. Given that the byproduct of extracting Cores from active compounds is the side chain fragments, it is natural to consider using side chains from two trees to build linkers. The linker construction process unfolds through three stages. First, two anchor atoms were chosen from two Core fragments as the linker growth anchors, named as **Growth Anchor** in Fig. 1C. Second, for each growth anchor, we extracted a side chain that grew from this anchor. Third, from each side chain, we selected an anchor atom to fuse them, designated as **Fusing Anchor** in Fig. 1C. By connecting two fusing anchors from distinct trees, core fragments from distinct targets can be consequently linked together. For growth anchors that have not been selected, decorating R-groups was considered to be helpful in improving the diversity of generated molecules. Side chains growing from these anchors can be sliced at different atoms to construct R-groups. Ultimately, as illustrated in Fig. 1C, the leaf node **Rationale** was constructed by decorating valid R-groups on unselected growth anchors. The final molecule was generated by fusing two rationales at their fusing anchors.

The sizes of the rationale libraries for GSK3*β*, JNK3, ROR*γ*t, and DHODH surpassed 130k, 60k, 150k, and 13k, respectively. It can be estimated that the chemical space accessible to AIx-Fuse ranges between 10^9^ and 10^10^ on both GSK3*β* | JNK3 and ROR*γ*t| DHODH benchmark. Virtual screening within chemical spaces of such magnitude demands an overwhelming amount of computational resources, often beyond reach. This challenge mirrors that encountered by AlphaZero when evaluating the value of massive states within the game space. Therefore, we did not assess the entire chemical space but instead employed iterative simulation and exploration by Monte Carlo tree search (MCTS). To guide the exploration, we also trained a multi-task graph neural network (GNN) to predict the docking scores against distinct targets, which is described in the next subsection.

### Graph Neural Network for Dual-Target Docking Score Prediction

We trained a multi-task GNN as a dual-target docking score prediction. Prior to the graph encoding by the GNN, the extraction of atomic features and bond features is imperative. We utilized a combination of nine atomic features and four bond features (refer to *Supplementary Table S3*), which were adopted by Xiong et al(42).

Graff et al. demonstrated that in the realm of molecular docking active learning, message-passing neural networks (MPNN) outperformed both Random Forest and traditional Neural Network approaches (35). Nonetheless, MPNN, when applied to the processing of molecular graphs, may encounter the issue of diluted effects. This arises from its uniform treatment of all nodes regardless of their distance to a target node, leading to weakened impacts of topologically adjacent nodes and functional groups (42). To address this limitation, attention mechanisms have been introduced, offering improved neighbor aggregation in the form of the Graph Attention Network (GAT) (62). More recently, the Attentive Fingerprints (Attentive FP) framework (42) was proposed as an approach capable of capturing both local atomic interactions through node information propagation and non-local effects within a molecule through a graph attention mechanism. It was tailored for molecular feature extractions and outperformed other architectures (including MPNN) across comprehensive benchmarks.

In our study, we applied the Attentive FP framework to the domain of molecular docking active learning. Two convolutional layers were constructed to extract atomic features, and a readout layer was utilized for generating molecular embeddings. Subsequently, docking scores were predicted by feeding molecular embeddings into a multi-task fully connected layer. The network architecture can be represented as Eq. 1, Eq. 2, and Eq. 3:

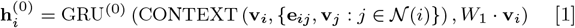

where **v**_*i*_ represents the raw node feature of vertex *i*, **v**_*j*_ is that of neighboring node *j*, **e**_*ij*_ denotes the raw feature of edge *ij, W*_1_ is a trainable weight matrix, and 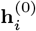 is the initial embedding of node *i*. The GRU used here is a variant of a long short-term memory (LSTM)(63) recurrent network unit, which has demonstrated effective information retention and filtering capabilities through the use of simplified update and reset gates.(64). The CONTEXT function used here served as an edge-level attentive aggregation.

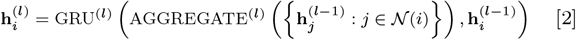

where 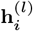 denotes the updated representation of atom *i* at layer *l*, and the AGGREGATE function used here served as a node-level attentive aggregation.

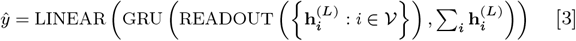

where 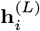 is the updated representation of atom *i* at the last layer, the READOUT function used here served as a graph-level attentive aggregation to extract the molecular embedding, and *ŷ* denotes the predicted values obtained by passing the molecular embedding through a fully connected layer.

The attention mechanism utilized in CONTEXT, AGGREGATE, and READOUT functions can be detailed described as Eq. 4, Eq. 6, and Eq. 8:

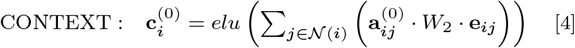

where

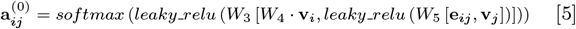

where *W*_2_, *W*_3_, *W*_4_, *W*_5_ are trainable weight matrices, leaky relu (65) and elu(66) are variations of the relu (67) nonlinear activation function, the activated results of leaky relu are further normalized using the softmax function to obtain the attentive weights 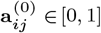, and the output 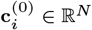 is the context messages activated by elu function.

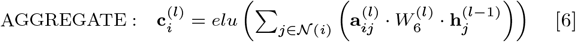

where

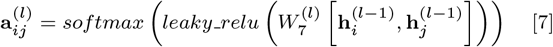

where *l >* 0 is the number of this layer, 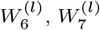 are trainable weight matrices at this layer, the attentive weights 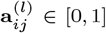 is used to aggregate the neighboring messages of node *i*, and the aggregation result 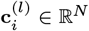 is to be processed by GRU and then be fed into next layer.

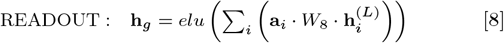

where

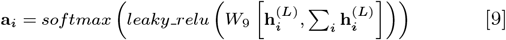

where *L* represents the final layer, *W*_8_, *W*_9_ are trainable weight matrices, the attentive weights **a**_*i*_ is used for aggregating the atomic representation, and the output **h**_*g*_ *∈*ℝ^*N*^ is the final representation at the molecular level.

We used MSELoss to measure the mean-squared error of our docking score regression model, which can be mathematically represented as:

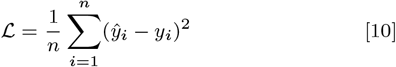

where *n* is the batch size, *y*_*i*_ represents the ground-truth docking score of *i*_*th*_ sample, and *ŷ*_*i*_ is the corresponding predicted value.

The entire network was trained in a multi-target fashion, yielding two predicted docking scores for a given molecule on two targets. During each training iteration, the dataset was randomly divided into a training dataset and a validation dataset in a 9:1 ratio. To mitigate the risk of overfitting, early stopping techniques were implemented. Specifically, we monitored each model’s performance on the validation set and continued training until no improvement was observed over 20 epochs or the maximum epochs of 1000 was reached.

Our models were trained on Pytorch(68) framework using the Adam(69) optimizer for gradient descent optimization. The Deep Graph Library (DGL) package(70) and the DGLLifeSci (71) extension were employed for the implementation of Attentive FP. All experiments were conducted within a uniform computational environment, consisting of 4 GPUs of Quadro RTX 5000, 96 CPU cores of Intel(R) Xeon(R) Gold 6248R, and the Ubuntu 20.04.6 LTS operating system. *Supplementary Text 5* describes the details of training and evaluation.

### Collaborative Learning of Pharmacophore Fusion and Molecular Docking

AIxFuse employs collaborative RL and AL to learn pharmacophore fusion patterns that satisfy the dual-target structural constraints simulated by molecular docking. As is shown in Fig. 1C, the RL framework is Actor-Critic-like, involving two MCTS Actors to generate pharmacophore-fused molecules and a dual-target docking score Critic trained by Active Learning. Algorithm 1 describes the collaborative learning procedure, where the INIT GEN function (i.e. AIxFuse (w/o RLAL)) generates the pharmacophore-fused molecules 𝒢 ^(0)^ by random sampling and fusing rationales from distinct trees, the DOCKING function dock generated molecules into target binding pockets to estimate their dual-target binding affinities, the TRAIN function represents active learning of the dual-target docking scores by a multi-task AttentiveFP model 𝒩, the trained parameters 𝒲 ^(0)^ of 𝒩 is then used in SELF PLAY function to generate a new batch of pharmacophore-fused molecules 𝒢 ^(*i*)^.

The INIT GEN function in Algorithm 1 is described in *Supplementary Text 3*. In short, AIxFuse initializes the weight of

#### Algorithm 1: Collaborative Learning of Pharmacophore Fusion and Molecular Docking

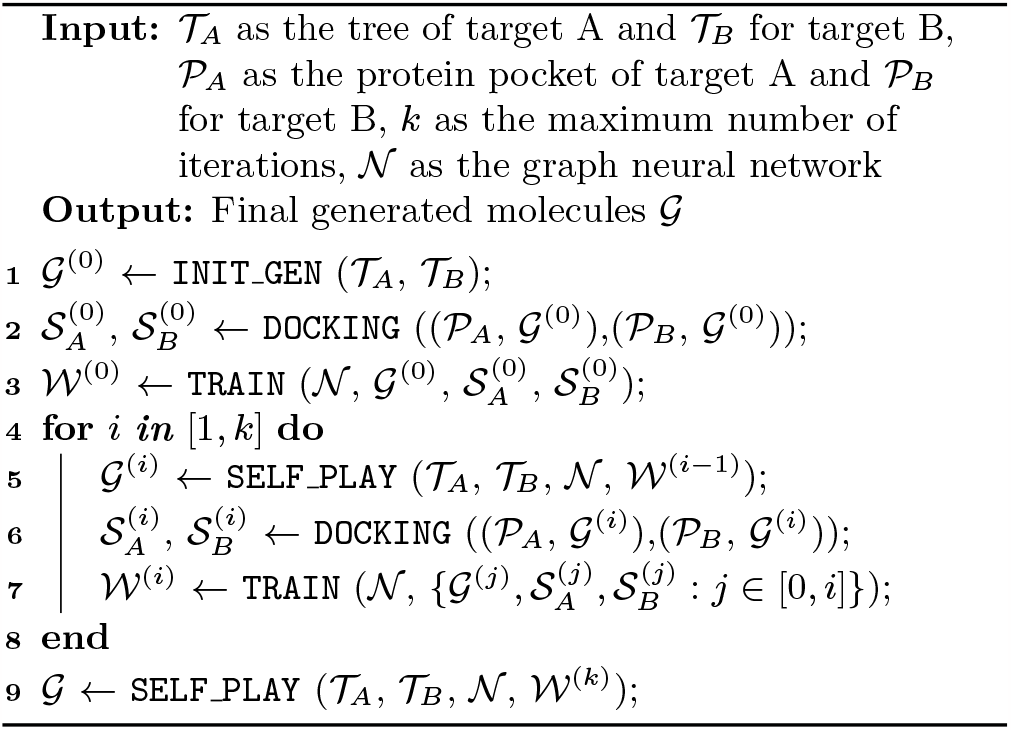

each node when building trees, and randomly samples child nodes according to their weights. The random-sampled rationales from two trees were fused to generate molecules.

The SELF PLAY function in Algorithm 1 is detailed in *Supplementary Algorithm 3*. Two MCTS Actors run in a self-paly manner, where we run simulations of pharmacophore fusion to reward each exploration. The reward function of a given molecule *m* can be formulated as:

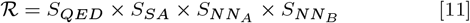

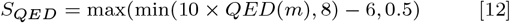

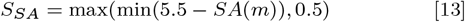

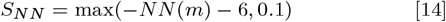

where *NN* predicts the docking score of *m* against target A (*NN*_*A*_) or target B (*NN*_*B*_), and the upper confidence bound (UCB) of reward expectation for a given node is used in the SELECT function of in *Supplementary Algorithm 3*, which can be represented as:

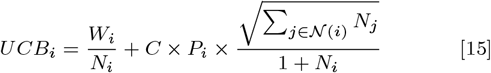

where *i* denotes the index of given node, *N*_*i*_ is the number of selection for this node, *W*_*i*_ represents the total reward of this node, *C* is a constant that controls the exploration tendency, *P*_*i*_ is the simulated reward of this node, and *j* is any brother node of node *i*.

### Computational Methods of MM/GBSA and ABFEP Calculations

Considering the huge computational consumption of molecular dynamics simulations, we conducted MM/GBSA binding free energy calculations for a subset of generated molecules. We initially selected the top 200 ranking molecules considering novelty, dual-target docking scores, 2D SNN, and 3D SNN. Subsequently, we filtered them by visual inspection, resulting in the retention of 134 molecules for MM/GBSA calculation. Among them, 17 representative molecules were selected for ABFEP calculation against ROR*γ*t and 7 out of 17 molecules were selected for ABFEP calculation against DHODH.

Molecular dynamic simulations were performed by GROMACS (version 2020.6)(72). AMBER99SB-ILDN(73) force field was used for proteins. General Amber Force Field (GAFF)(74) with AM1-BCC(75, 76) partial charges were used for ligands. The proteinligand complex was solvated in the dodecahedral box of TIP3P(77) water with the minimum distance between the solute and the box of 10 Å. The system was minimized using the steepest descent algorithm with max steps of 10,000, followed by 100 ps NVT equilibration at 300 K and then 100 ps NPT equilibration at 300 K and 1 bar. Subsequently, 2 ns NPT production was performed at 300 K and 1 bar and the frames were saved every 10 ps for further analysis and MMGBSA calculations. The distance cutoff of short-range interactions was set to 10 Å. Long-range electrostatic interactions were treated with PME algorithm(78). LINCS(79) algorithm was applied to constrain the hydrogen-involved bonds and all the MD simulations were performed with a timestep of 2 fs. The MM/GBSA calculations were performed with AMBER(80) with a time interval of 20 ps for extracted frames via the singletrajectory scheme, where the structures of the separate protein and ligand were extracted from the trajectory of the complex. ABFEP calculations were performed following the schemes employed by Aldeghi et al (81, 82).

## Data Availability

Data and code will be soon available at https://github.com/biomed-AI.

## ACKNOWLEDGMENTS

This work has been supported by the National Key Research and Development Program of China (2020YFB0204803) and the Guangzhou Science and Technology Research Plan (202007030010).

