## Supporting Information for "Structure-Aware Dual-Target Drug Design through Collaborative Learning of Pharmacophore Combination and Molecular Simulation"

**This PDF file includes:**

Supporting text
Figs. S1 to S6
Tables S1 to S6
SI References

### Supporting Information Text

### 1. Data process

For each target, we used the following process to collect and preprocess the active molecules. Firstly, active molecules against each target are collected from CHEMBL and BindingDB separately. Secondly, active molecules were filtered by IC50 (our tasks focus on inhibitor design) value, and only the compounds with IC50 lower than 1,000 nM were retained. Thirdly, we de-duplicate the remaining compounds by canonical SMILES. Finally, for each target, the number of non-redundant active molecules is shown in Supplementary Table S1. These molecules were used to construct the active molecule library as a reference.

In order to obtain the protein-ligand interactions between those molecules and target proteins, we need to dock them to obtain complex structures. Therefore, it is necessary to prepare the protein acceptor pocket. For ROR$\gamma$t and DHODH targets, referring to the work of Chen et al.(1), we used the protein co-crystal structures with PDB id 5NTP(2) and 6QU7(3). We removed other chains in the original structures and performed hydrogen addition, hydrogen bond assignment, water removal, restrain minimization, etc. The above operations were performed in protein prepare(4), using default parameters. The docking grid was centered according to the ligand position, and the bounding box was set to 15 Å. For small molecules, we ran ligand preparation(5) using Epik to set the ionization possible state under neutral conditions and generated up to 32 possible chiral isomers per molecule. Then we used Glide for docking, using XP precision (consistent with the work of Chen et al.) and the default value for other parameters. For the GSK3$\beta$ and JNK3 targets, we used the protein co-crystal structures with PDB ids 6Y9S(6) and 4WHZ(7) as raw data. We repeated the above steps in ROR$\gamma$t|DHODH preparation for GSK3$\beta$| JNK3 task, but we used the more common SP precision when docking.

### 2. Core Fragment Extraction and Scoring

Here are some detailed descriptions of core fragment extraction and scoring in our method. The overall extraction and selection process is outlined in Supplementary Algorithm 1, where the PLIP function employs PLIP(8, 9) to extract PLIs from complex structures, COMB enumerates valid combinations of PLIs, the EXTR function is detailed in Supplementary Algorithm 2, and SCORE_SIZE, SCORE_INTER, and SCORE_DIS assign scores to the substructures. EXTR takes a given molecule and pharmacophore as input and extracts the minimal substructure that preserves this pharmacophore. The pharmacophore is defined as the combination of protein-ligand interactions (PLIs).

**Algorithm 1:** Extracting and selecting core fragments from active compounds

**Input:** Active molecules $\mathcal{M}$, protein-ligand complexes $\mathcal{C}$ and key residues $\mathcal{R}$
**Output:** Top-scoring core fragments $\mathcal{F}$

$\mathcal{D}_F \leftarrow$ empty dictionary `// Accumulating score for each fragment` ;
**for** $m$, $c$ ***in*** $\mathcal{M}$, $\mathcal{C}$ **do**
  $\mathcal{I} \leftarrow$ PLIP($c$) `// PLIs`;
  $\mathcal{P} \leftarrow$ COMB($\mathcal{I}$) `// Phamacophores as combinations of PLIs`;
  $\mathcal{M}_F \leftarrow$ empty dictionary `// Recording the max score for each fragment`;
  **for** $p$ ***in*** $\mathcal{P}$ **do**
    $f \leftarrow$ EXTR($m$,$p$) `// Core fragments`;
    $s_s \leftarrow$ SCORE_SIZE($f$,$m$);
    $s_i \leftarrow$ SCORE_INTER($f$,$\mathcal{I}$);
    $s_d \leftarrow$ SCORE_DIS($f$,$c$,$\mathcal{R}$);
    $s \leftarrow s_s \times (s_i + s_d)$ `// Scores`;
    $m_f \leftarrow$ MAX($m_f$, $s$) `//` $m_f \in \mathcal{M}_F$;
  **end**
  **for** $m_f$ ***in*** $\mathcal{M}_F$ **do**
    $d_f \leftarrow d_f + m_f$ `//` $d_f \in \mathcal{D}_F$;
  **end**
**end**
$\mathcal{F} \leftarrow$ Top-scoring $f$ in $\mathcal{D}_F$;

**Algorithm 2:** Core Fragment Extraction with Given Pharmacophore

**Input:** $m$ as given molecule, $p$ as given pharmacophore
**Output:** core fragment $f$

**Function** `EXTR`($m$,$p$):
  $\mathcal{L} \leftarrow$ empty set `// set of interacted atoms`;
  **for** $i$ ***in*** $p$ **do**
    $\mathcal{L}$.add($a$) **for** $a$ **in** GET_ATOMS($m$,$i$) `// get interacted atoms`;
  **end**
  $\mathcal{S} \leftarrow$ empty set `// set of minimum substructure atoms`;
  **for** $a$ ***in*** $\mathcal{L}$ **do**
    **for** $b$ ***in*** $\mathcal{L}$ **do**
      $\mathcal{P} \leftarrow m$.findShortestPath($a$,$b$) `// get atoms in the shortest path`;
      $\mathcal{A} \leftarrow \mathcal{P}$.extendRingAtoms `// expand ring atoms in the path`;
      $\mathcal{S}$.add($a$') **for** $a$' **in** $\mathcal{A}$ `// get minimum substructure atoms`;
    **end**
  **end**
  $f \leftarrow m$.getSubstructure($\mathcal{S}$) `// get minimum substructure`;
  **return** $f$;

The SCORE_SIZE function, SCORE_INTER function, and SCORE_DIS function were used as scoring functions of the extracted core fragment. They can be formulated as below:

SCORE_SIZE rewards fragment that is neither too small nor too large, which is formulated by Eq. 1:

$$s_s = \begin{cases} 0 & h < \frac{N}{6} || h > \frac{5\times N}{6} \\ 1 & \frac{N}{3} < h\frac{2\times N}{3} \\ \frac{6\times h}{N} - 1 & \frac{N}{6} \leq h \leq \frac{N}{3} \\ -\frac{6\times h}{N} + 5 & \frac{2\times N}{3} \leq h \leq \frac{5\times N}{6} \end{cases} \quad [1]$$

, where h is the number of heavy atoms in the fragment and N represents the number of heavy atoms in the source molecule.

SCORE_INTER rewards fragment that shares more PLIs, and is represented as Eq. 2:

$$s_i = \sum_{i\in\mathcal{I}} \sum_{a\in i\&f} 1/len(i) \quad [2]$$

, where $i$ enumerates each PLI in $\mathcal{I}$, a is any atom in core fragment $f$ that contributes to PLI $i$, $len(i)$ counted the number of ligand atoms that share this PLI.

SCORE_DIS rewards fragment that is closer to given key residues, which is defined as the sum of the distance score obtained by each atom in this fragment:

$$s_d = \sum_{a\in f} s_a \quad [3]$$

, where the distance score $s_a$ of each atom can be expressed as a piecewise function:

$$s_a = \begin{cases} 0 & d_a > 6 \\ 1 & d_a < 3 \\ -\frac{d_a}{3} + 2 & 3 \leq d_a \leq 6 \end{cases} \quad [4]$$

, where the minimum distance $d_a$ between a given atom and key residues is calculated by:

$$d_a = \min_{b\in\mathcal{K}} dis(a, b) \quad [5]$$

, where $b$ Enumerates key interacting residues in $\mathcal{K}$.

### 3. Tree Initialization for Pharmacophore Fusion

In the INIT_GEN function of Algorithm 1 in the main text, AIxFuse iteratively and randomly samples child nodes in the trees until the leaf node of rationale and then fuse them to generate molecules. The sampling probability of each child node is determined by its weight that is initialized when building trees:

$$\mathcal{P}_n = \frac{\mathcal{W}_n}{\sum_{b\in\mathcal{B}_n} \mathcal{W}_b} \quad [6]$$

, where $\mathcal{B}_n$ is the set of brother nodes of node $n$. The weights of nodes in each level are initialized when building trees. For each Core, we accumulate the negative base 10 logarithms of the I/EC50 values of the active molecules that share this Core as its node weights. For each Growth Anchor, we accumulate the heavy atom number (8 if $> 8$) of all its Side Chains as its node weight. Since a Fusing Anchor may be shared by multiple active molecules, we accumulate the number of active molecules that share this Fusing Anchor as its node weight. The rationale is determined by the combination of Fusing Anchor and R-groups. The R-groups are the combination of Fusing Anchors from other Growth Anchors. Therefore, the weight of Rationale is the cumulative multiplication of the weights of these Fusing Anchors.

### 4. Implement of Other Methods

For the GSK3$\beta$|JNK3 benchmark task, both RationaleRL and MARS utilized activity predictors developed by Li et al. (10) to estimate molecular activity. We aligned REINVENT2.0 with these approaches. We ran RationaleRL and MARS by the tutorials in their GitHub pages[*†]. For RationaleRL we directly ran *python decode.py −−model ckpt/gsk3_jnk3_qed_sa/model.final −−num_decode 500 > outputs.txt* and generated 18,500 molecules. We then randomly sampled 10,000 molecules as the generation result of RationaleRL. For MARS we ran *python -m MARS.main −−train −−run_dir runs/gsk3b_jnk3 −−num_mols 10000* and employed the molecules of converged step (651) as the generation result of MARS. For REINVENT2.0 we changed the scoring_function configuration in the Reinforcement_Learning_Demo.ipynb of ReinventCommunity[‡]. We replaced the original properties with qed_score, SA_score, and the predicted GSK3$\beta$ and JNK3 activity scores that are used in RationaleRL and MARS. After that, we first ran Reinforcement_Learning_Demo.ipynb to train the agent and then ran Sampling_Demo.ipynb to generate 10,000 molecules.

For the ROR$\gamma$t|DHODH benchmark task, the machine learning models for molecular activities evaluation have not been established before. Here we trained two Support Vector Machine (SVM) models for ROR$\gamma$t and DHODH activity prediction, respectively. However, the training procedure like Li et al.(10) is not feasible. Because there are too few (113) DHODH inactive molecules in ExCAPE-DB to train the machine-learning model. Here we assumed that most molecules in CHEMBL are not active to DHODH if their assay information is not related to DHODH. We first collected molecules from CHEMBL and curated an “inactive” molecule dataset that is 5 times larger than the active dataset (the size of the active dataset is shown in Supplementary Table S1). We then split the dataset into a training set and a test set according to the molecular similarity. Each molecule in the test set was lower than 0.8 Tanimoto similarity with any molecules in the training data. The size of training data and test data is shown in Supplementary Table S2. We used the molecular fingerprint features of 2048 dimensions as input. The hyperparameters of the SVM models are optimized by Bayesian optimization. As is shown in Supplementary Fig. S1, the Area Under ROC Curve (AUC) obtained on DHODH and ROR$\gamma$t were 0.998 and 0.996, respectively. We then used these two activity predictors in RationaleRL, MARS, and REINVENT2.0 to generate molecules for ROR$\gamma$t|DHODH benchmark task.

### 5. Implementation of GNN training and evaluation

Supplementary Table S3 shows the molecular representation (11) used by our graph neural network. Most of these features are encoded by one-hot representation, except for formal charge and radical electron number which are represented as integers due to their additive properties. To construct a one-hot encoding feature, all possible categorical variables related to the feature are listed and assigned values of either 1 or 0 (one-hot or null) based on their correspondence to those variables. For instance, a 16-bit vector is designated for encoding atomic symbols, while a 6-bit vector is employed for encoding hybridization states. Notably, atomic chirality is encoded using three distinct bits: one indicating the presence of a chiral center, and the remaining two bits specifying whether it is in R-form or S-form. Additionally, stereotypes of double bonds are denoted by a feature that distinguishes their potential E/Z configurations.

The training of our graph neural network went through 5 iterations. In the first round of training, 10,000 molecules randomly generated by INIT_GEN were used for model training. After the first round of training is completed, the obtained model was used for the reward function of the second round of Monte Carlo tree search. In the generation procedure of the second round, AIxFuse generated 10,000 molecules that were non-redundant with previous molecules. We then integrated all generated molecules as a new training dataset and the second round of model training was performed to obtain a new docking scoring prediction model. Iteratively, the fifth round of model training is finally completed. Using the model obtained from the fifth round, AIxFuse generated 10,000 molecules as its final result.

To evaluate the performance of the model at each round, we constructed a test set based on the final generated molecules. The molecules in the test set were de-redundant with all molecules in the training set. We used the model trained for each epoch to make predictions on the training set. The performance evaluation metrics included MSE and $R^2$. The formula of MSE is as follows:

$$MSE = \frac{1}{n}\sum_{i=1}^{n}(y_i - \hat{y}_i)^2 \quad [7]$$

, where $y_i$ represents the ground-truth value, $\hat{y}_i$ represents the predicted value of the model, and $n$ represents the sample size. The formula squares the difference between each observed value and the corresponding model-predicted value and averages it

[*] https://github.com/wengong-jin/multiobj-rationale
[†] https://github.com/bytedance/markov-molecular-sampling
[‡] https:// github.com/MolecularAI/ReinventCommunity

over all samples. The formula of $R^2$ is as follows:

$$R^2 = 1 - \frac{\sum_{i=1}^{n}(y_i - \hat{y}_i)^2}{\sum_{i=1}^{n}(y_i - \bar{y})^2} \tag{8}$$

, where $\bar{y}$ is the average of ground-truth value.

### 6. Generating Pharmacophore-Fused Molecules by two Self-Play MCTS Actors

Supplementary Algorithm 3 describes the SELF_PLAY function in Algorithm 1 of the main text. Here the REWARD function $\mathcal{R}$ of any generated or simulated molecule $m$ can be formulated as:

$$\mathcal{R} = S_{QED} \times S_{SA} \times S_{NN_A} \times S_{NN_B} \tag{9}$$

$$S_{QED} = \max(\min(10 \times QED(m), 8) - 6, 0.5) \tag{10}$$

$$S_{SA} = \max(\min(5.5 - SA(m)), 0.5) \tag{11}$$

$$S_{NN} = \max(-NN(m) - 6, 0.1) \tag{12}$$

, where $NN$ predicts the docking score of $m$ against target A ($NN_A$) or target B ($NN_B$), and the upper confidence bound (UCB) of reward expectation for a given node is used in the SELECT function of MCTS, which can be represented as:

$$UCB_i = \frac{W_i}{N_i} + C \times P_i \times \frac{\sqrt{\sum_{j \in \mathcal{N}(i)} N_j}}{1 + N_i} \tag{13}$$

, where $i$ denotes the index of given node, $N_i$ is the number of selection for this node, $W_i$ represents the total reward of this node, $C$ is a constant that controls the exploration tendency, $P_i$ is the simulated reward of this node, and $j$ is any brother node of node $i$.

**Algorithm 3:** SELF_PLAY to Generate Pharmacophore-Fused Molecules

**Input:** $\mathcal{T}_A$ as tree of target A and $\mathcal{T}_B$ for target B, $\mathcal{N}$ as the neural network, and $\mathcal{W}$ as its weight parameters, and n as the number of generated molecules

**Output:** Generated molecules $\mathcal{G}$

$\mathcal{G} \leftarrow$ empty list `// list of generated molecules`;
**while** $LEN(\mathcal{G}) < n$ **do**
    $\mathcal{C}_A$, $\mathcal{C}_B \leftarrow \mathcal{T}_A$.root.children,$\mathcal{T}_B$.root.children `// Cores`;
    $\mathcal{V}_A$, $\mathcal{V}_B \leftarrow$ `SELECT` ($\mathcal{C}_A$), `SELECT` ($\mathcal{C}_B$) `// best Core`;
    $\mathcal{S} \leftarrow$ empty list `// list to record the simulation results`;
    $\mathcal{S}$, $m \leftarrow$ `SIMULATE` ($\mathcal{S}$, $\mathcal{V}_A$, $\mathcal{V}_B$, $\mathcal{N}$, $\mathcal{W}$);
    $\mathcal{G}$.append($m$) `// record` $m$ `as generated molecules`;
    **if** *Not FINAL_GEN* **then**
        $\mathcal{G}$.extend(`SAMPLE` ($\mathcal{S}$)) `// to improve the training set diversity`;
    **end**
**end**
**Function** `SIMULATE`($\mathcal{S}$, $\mathcal{V}_A$, $\mathcal{V}_B$, $\mathcal{N}$, $\mathcal{W}$):
    $\mathcal{C}_A$, $\mathcal{C}_B \leftarrow \mathcal{V}_A$.children, $\mathcal{V}_B$.children;
    $\mathcal{M}_A$, $\mathcal{M}_B \leftarrow$ `SAMP_MOL` ($\mathcal{C}_A$, $\mathcal{C}_B$) `// sample Rationales to fuse molecules`;
    $\mathcal{S}$.extend($\{\mathcal{M}_A, \mathcal{M}_B\}$) `// record simulation results`;
    `UPDATE_P` ($\mathcal{C}_A$, $\mathcal{C}_B$, `REWARD` ($\{\mathcal{M}_A, \mathcal{M}_B\}$),$\mathcal{N}$, $\mathcal{W}$) `// update P in Eq.13`;
    $\mathcal{V}'_A$, $\mathcal{V}'_B \leftarrow$ `SELECT` ($\mathcal{C}_A$), `SELECT` ($\mathcal{C}_B$) `// best child`;
    **if** `IS_INSTANCE` *($\mathcal{V}'_A$, Rationale)* **then**
        $m \leftarrow$ `FUSE` ($\mathcal{V}'_A$, $\mathcal{V}'_B$) `// fuse Rationales to generate molecules`;
        `UPDATE_W_N` ($\mathcal{V}'_A$, $\mathcal{V}'_B$, `REWARD` ($m$, $\mathcal{N}$, $\mathcal{W}$)) `// update N, W in Eq.13`;
        **return** $\mathcal{S}$, $m$
    **end**
    **return** `SIMULATE` ($\mathcal{S}$, $\mathcal{V}'_A$, $\mathcal{V}'_B$, $\mathcal{N}$, $\mathcal{W}$) `// recursive call`;

### 7. Supplementary Results

In Table 1 of the main text, we report the similarity of the generated molecules to the active molecules. Here we are also interested in understanding the distance between the property distribution of generated molecules and active molecules. As is shown in Table S4, Fréchet ChemNet Distance (FCD)(12) and Similarity to Nearest Neighbor (SNN) are both used to evaluate the molecular similarity between generated compounds and inhibitors of GSK3$\beta$, JNK3, ROR$\gamma$t, and DHODH. When compared with REINVENT2.0, RationaleRL, and MARS, AIxFuse achieved the minimum FCD on all four targets. This is reasonable since molecules generated in the chemical space gaps between inhibitors of two targets are supposed to be similar to both ends. For SNN we found that AIxFuse obtained the highest similarity to inhibitor libraries except ROR$\gamma$t's. But when we studied the property distribution distance to ROR$\gamma$t inhibitors, AIxFuse generated compounds achieved minimum distance for QED, SA, LogP, and molecular weight, which is consistent with FCD. This is probably because, for molecular generation, FCD is more suitable than SNN for similarity evaluation (12). Ranking property distance to GSK3$\beta$ and JNK3 inhibitors ascendingly, AIxFuse was $1_{st}|1_{st}$, $2_{nd}|2_{nd}$, $1_{st}|2_{nd}$ and $3_{rd}|1_{st}$ on QED, SA, LogP and molecular weight, respectively. For ROR$\gamma$t and DHODH, AIxFuse achieved minimum distribution distance on most properties except molecular weight. The detailed density distributions are visualized in Supplementary Fig. S2.

In Figure 4C of the main text, we plotted the distribution of dual-target docking scores of generated molecules. We found that besides AIxFuse, RationaleRL also successfully generated some molecules that obtained better docking scores than **(R)-14d**. However, further study found that these molecules contain a large number of benzene ring fragments (examples shown in Supplementary Fig. S3A). Due to the hydrophobic nature of the benzene ring, the LogP of RationaleRL-generated molecules is concentrated above 7 (shown in Supplementary Fig. S2G). This could potentially lead to the problem of insufficient solubility. MARS generated very few molecules with docking scores lower than **(R)-14d**. This could be due to that its generated molecules are too small (illustrated in Supplementary Fig. S2H) to satisfy the binding modes required by both targets simultaneously.

### 8. Visual Inspection

We comprehensively considered the dual-target Docking scoring of the generated molecules, as well as the 2D SNN and 3D SNN to the active molecules of different targets, and selected the top 200 molecules from the 10,000 generated by AIxFuse. For the top 200 molecules, we use manual visual inspection to filter out molecules with unreasonable Docking poses. For example, we observed that in the molecular docking pose of some generated molecules, the PLI pattern between the core fragments and proteins has changed. It became different from the corresponding single-target active molecules.

Supplementary Fig. S4 shows a more intuitive example. As shown in Supplementary Fig. S4A, the generated molecule $M$ consists of red core fragments from active compounds of ROR$\gamma$t and blue from DHODH, which are called Fragment $F_1$ and Fragment $F_2$, respectively. Supplementary Fig. S4B visualizes a positive compound of target A, which also consists of fragment $F_1$. In the docking pose of the ROR$\gamma$t active compound, fragment $F_1$ interacts with residue 479 of target A, as shown in Supplementary Fig. S4D. However, in the low-energy docking pose of molecule $M$ in Supplementary Fig. S4C, fragment $F_1$ does not interact with HIE479. Therefore, for molecules with similar problems, we filter them out by visual inspection. In the end, we obtained 136 molecules that passed the filter and performed MM/GBSA calculations on them.

Supplementary Fig. S5 supplemented the ROR$\gamma$t|DHODH results of Figure 3A-D in the main text. Supplementary Fig. S6 supplemented the ROR$\gamma$t|DHODH results of Figure 2E in the main text. Overall the results in ROR$\gamma$t|DHODH were consistent with results in GSK3$\beta$|JNK3 task. Supplementary Table S5 illustrated the SMILES of our drug candidates for MM/GBSA and ABFEP calculation, and Supplementary Table S6 showed the results.

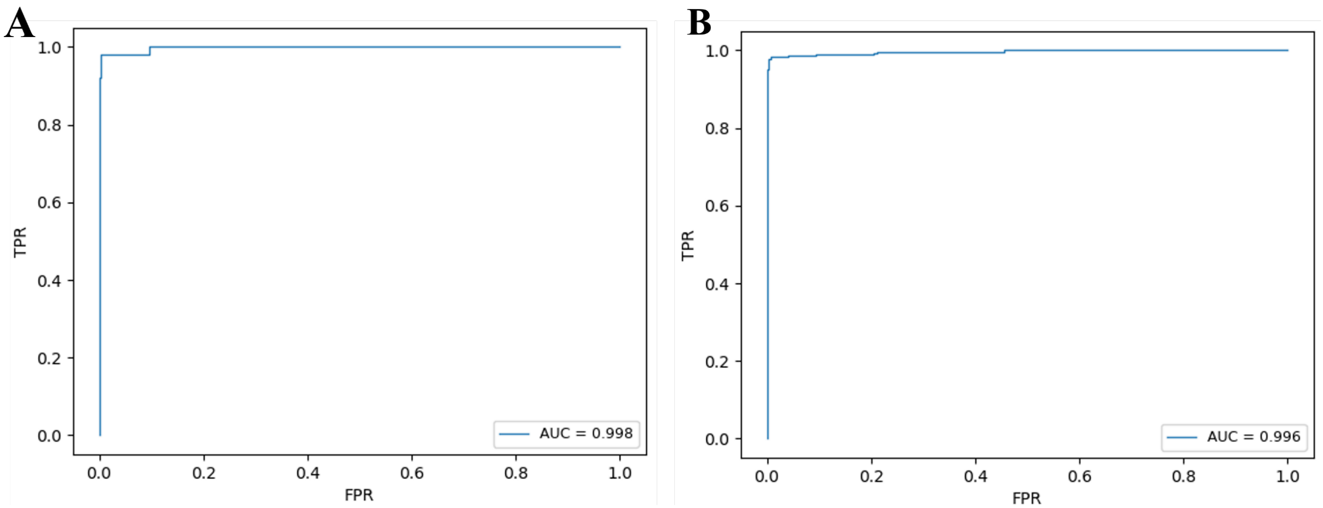

**Fig. S1.** ROC curves for the predictive model of A.DHODH and B.ROR$\gamma$t

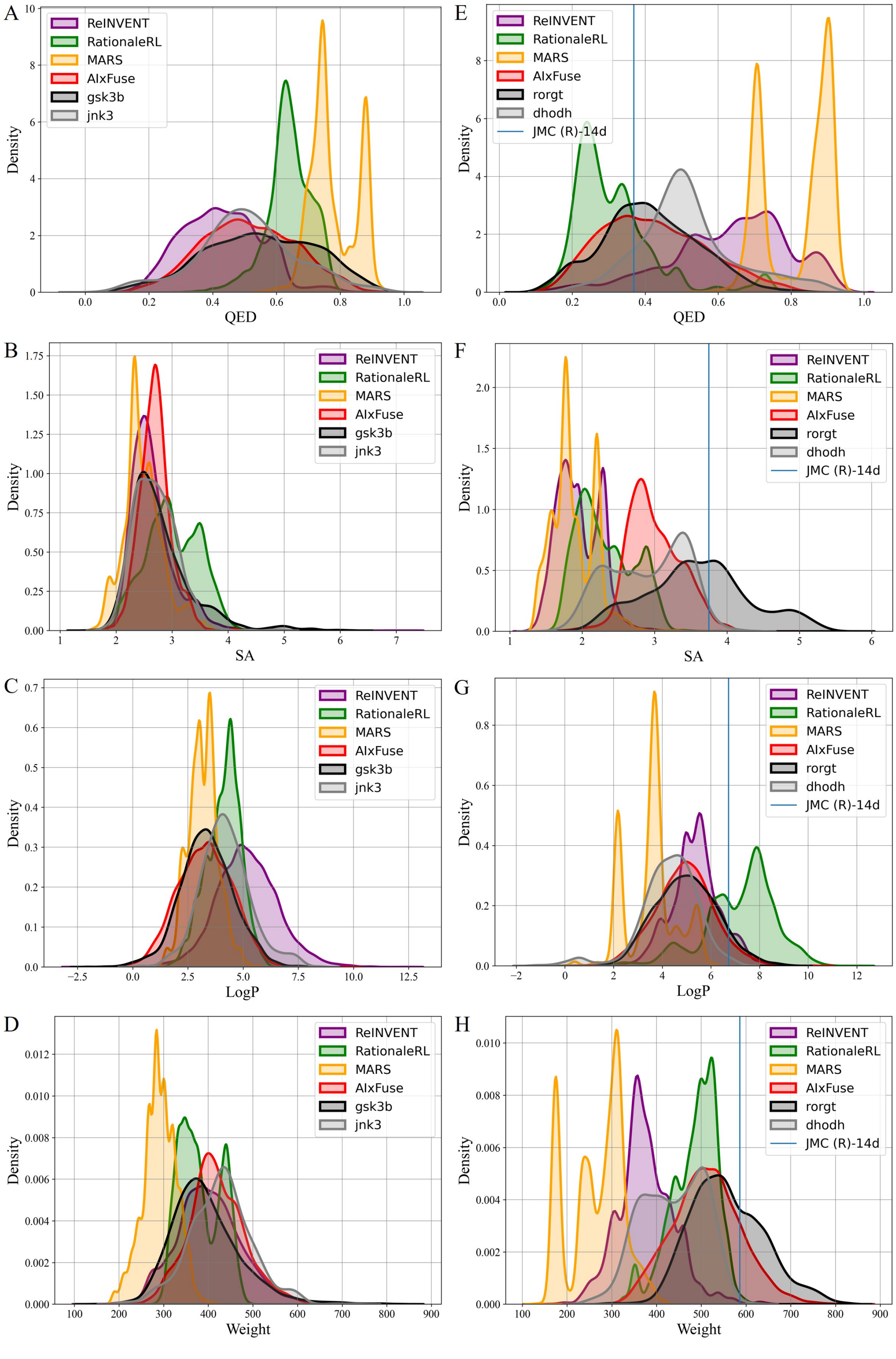

**Fig. S2.** The property distribution of molecules generated for GSK3$\beta$|JNK3 (A: QED, B: SA, C: LogP, D: Weight) and property distribution of molecules generated for ROR$\gamma$t|DHODH (E: QED, F: SA, G: LogP, H: Weight)

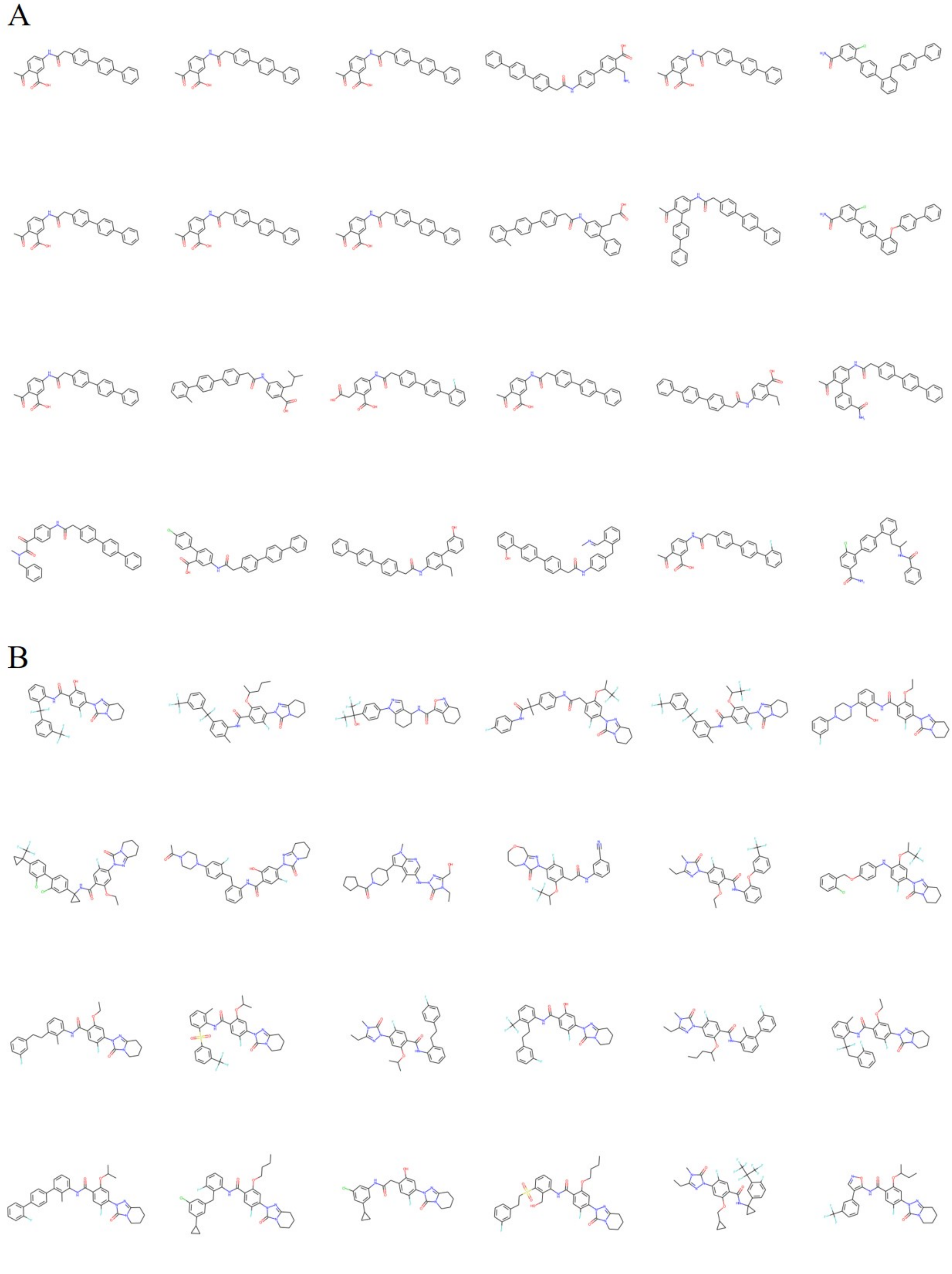

**Fig. S3.** The examples molecules generated by RationaleRL (A) and AIxFuse(B) that obtained better docking score than **(R)-14d** on ROR$\gamma$t

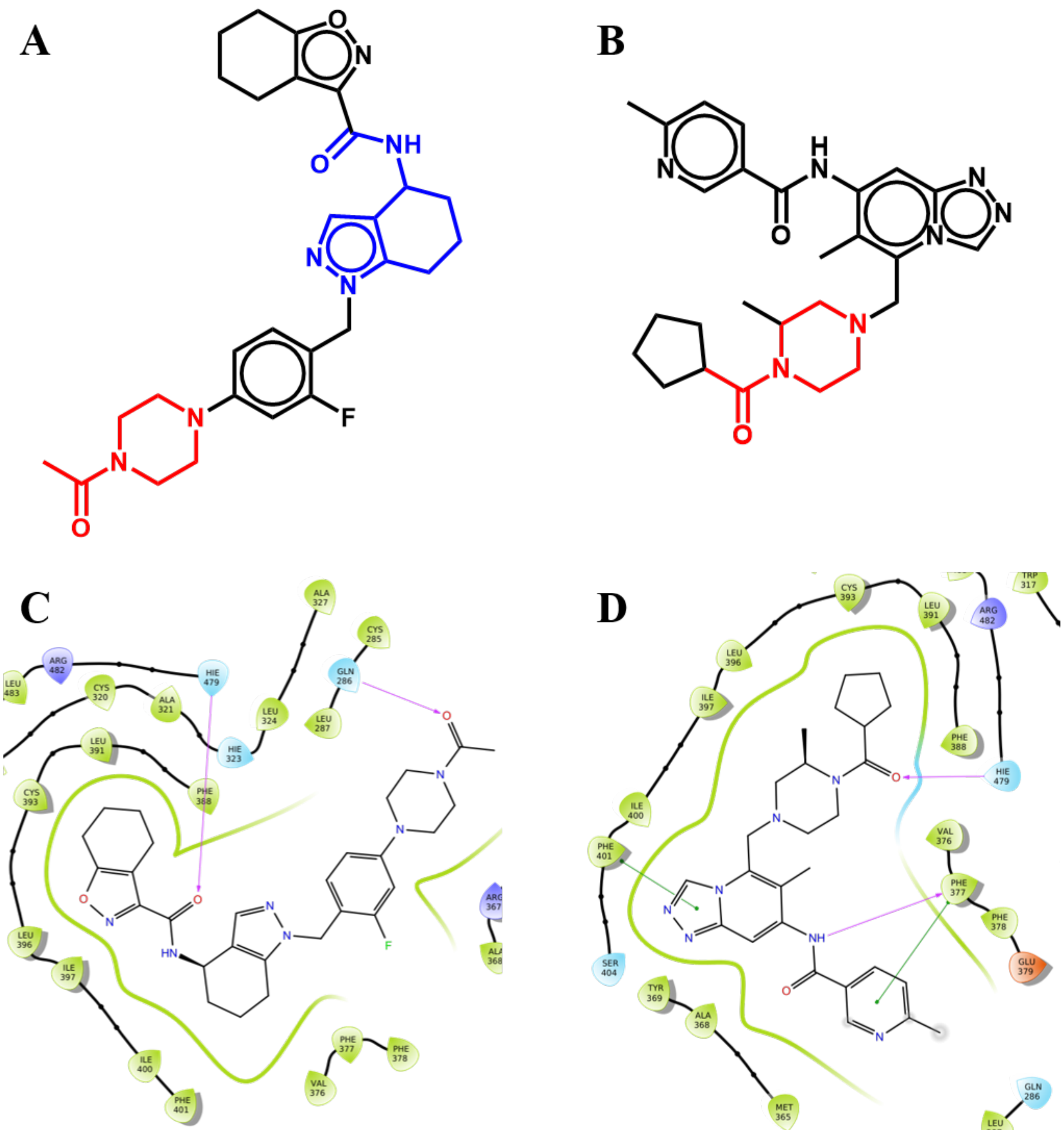

**Fig. S4.** An example of visual inspection: (A) A generated molecule $M$ with red core fragment $F_1$ from ROR$\gamma$t active compounds and blue core fragment $F_2$ from DHODH. (B) A ROR$\gamma$t active compound with corresponding core fragment $F_1$. (C) The ROR$\gamma$t Docking pose of generated molecule $M$. (D) The Docking pose of the corresponding ROR$\gamma$t active compound

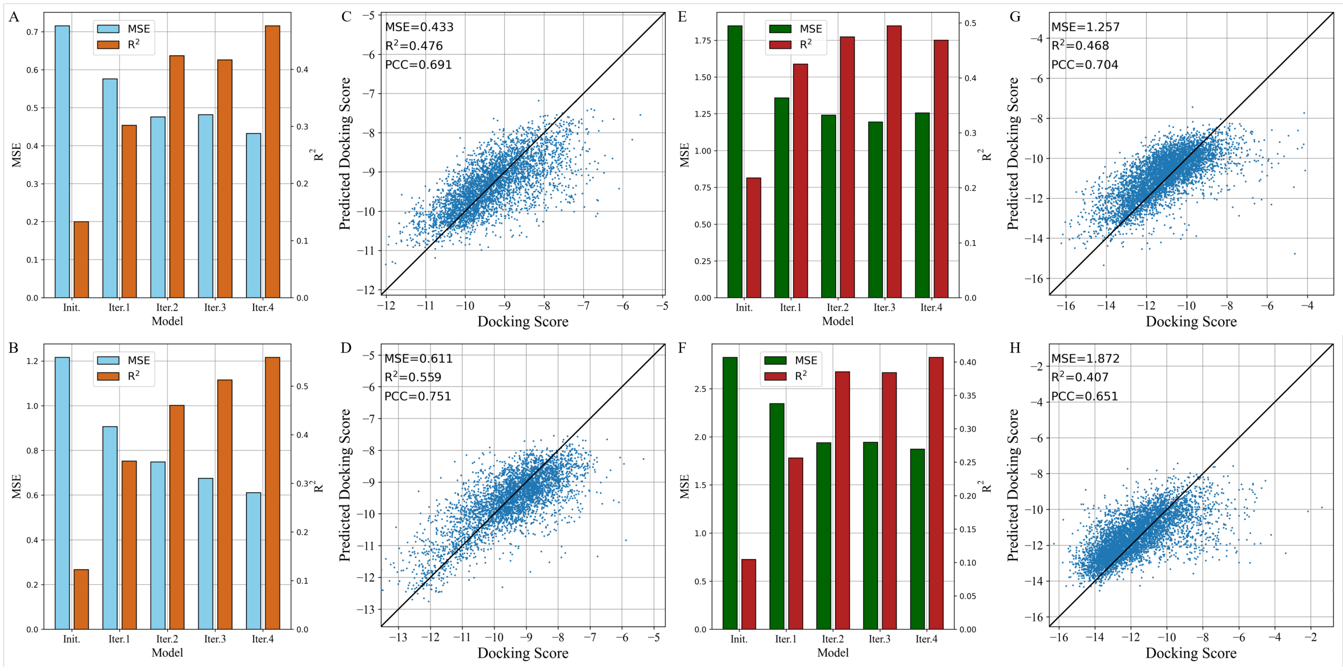

**Fig. S5.** Docking score prediction performance for (A) GSK3$\beta$, (B) JNK3, (E) ROR$\gamma$t, and (F) DHODH on different reinforcement learning iterations, and the scatter plot of predictive docking score of final model and the ground-truth value for (C) GSK3$\beta$, (D) JNK3, (G) ROR$\gamma$t and (H) DHODH

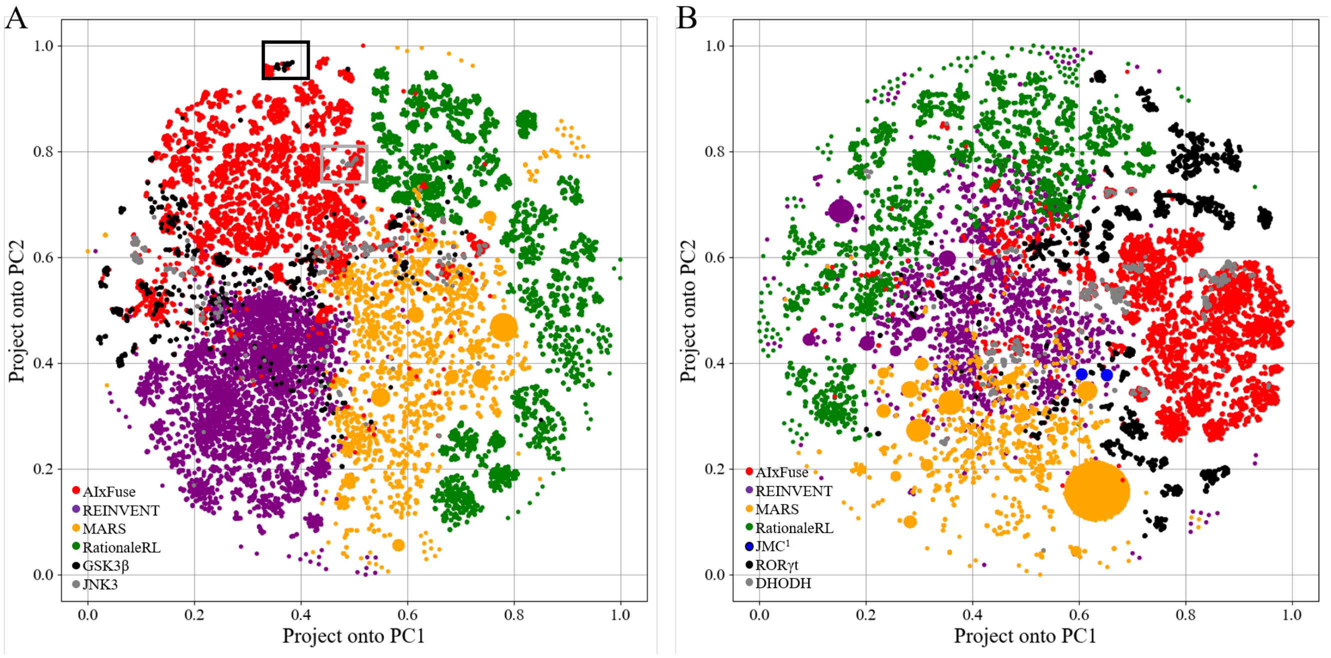

**Fig. S6.** The t-SNE visualization of AIxFuse's generated compounds on dual-target inhibitor design task for (A) GSK3$\beta$|JNK3 and (B) ROR$\gamma$t|DHODH.

**Table S1. Number of active compounds for different targets**

| Target | Number of Active Compounds |
|---|---|
| GSK3$\beta$ | 2128 |
| JNK3 | 791 |
| ROR$\gamma$t | 5902 |
| DHODH | 1210 |

**Table S2. Number of Training and Test dataset for ROR$\gamma$t and DHODH activity predictors.**

| Size of Dataset | Training | Test |
|---|---|---|
| ROR$\gamma$t | 10740 | 1686 |
| DHODH | 3876 | 300 |

**Table S3. Initial Atomic and Bond Features**

| atom feature | size | description |
|---|---|---|
| atom symbol | 16 | [B, C, N, O, F, Si, P, S, Cl, As, Se, Br, Te, I, At, metal] (one-hot) |
| degree | 6 | number of covalent bonds [0,1,2,3,4,5] (one-hot) |
| formal charge | 1 | electrical charge (integer) |
| radical electrons | 1 | number of radical electrons (integer) |
| hybridization | 6 | [sp, $sp^2$, $sp^3$, $sp^3d$, $sp^3d^2$, other] (one-hot) |
| aromaticity | 1 | whether the atom is part of an aromatic system [0/1] (one-hot) |
| hydrogens | 5 | number of connected hydrogens [0,1,2,3,4] (one-hot) |
| chirality | 1 | whether the atom is chiral center [0/1] (one-hot) |
| chirality type | 2 | [R, S] (one-hot) |
| **bond feature** | **size** | **description** |
| bond type | 4 | [single, double, triple, aromatic] (one-hot) |
| conjugation | 1 | whether the bond is conjugated [0/1] (one-hot) |
| ring | 1 | whether the bond is in ring [0/1] (one-hot) |
| stereo | 4 | [StereoNone, StereoAny, StereoZ, StereoE] (one-hot) |

**Table S4. Molecular similarities and property distribution distance between generated molecules and inhibitors of GSK3$\beta$, JNK3, ROR$\gamma$t, and DHODH**

| | GSK3$\beta$\|JNK3 | | | |
|---|---|---|---|---|
| **Method** | **REINVENT2.0** | **RationaleRL** | **MARS** | **AIxFuse** |
| **FCD** | 29.5\|30.8 | 28.2\|27.8 | 53.8\|46.8 | **16.7**\|**24.9** |
| **SNN** | 0.35\|0.41 | 0.38\|0.44 | 0.36\|0.45 | **0.47**\|**0.45** |
| **Dis. QED** | 0.12\|0.08 | 0.11\|0.14 | 0.23\|0.27 | **0.04**\|**0.02** |
| **Dis. SA** | **0.14**\|**0.07** | 0.28\|0.32 | 0.30\|0.21 | 0.18\|0.09 |
| **Dis. LogP** | 1.84\|0.99 | 0.80\|**0.26** | 0.43\|1.03 | **0.10**\|0.94 |
| **Dis. Weight** | **11.1**\|27.1 | 22.5\|49.9 | 103.7\|136.6 | 27.9\|**10.9** |
| | **ROR$\gamma$t\| DHODH** | | | |
| **Method** | **REINVENT2.0** | **RationaleRL** | **MARS** | **AIxFuse** |
| **FCD** | 39.1\|39.7 | 38.2\|35.6 | 59.2\|49.0 | **23.7**\|**14.1** |
| **SNN** | **0.45**\|0.37 | 0.44\|0.37 | 0.41\|0.38 | 0.31\|**0.60** |
| **Dis. QED** | 0.22\|0.12 | 0.10\|0.20 | 0.40\|0.30 | **0.02**\|**0.09** |
| **Dis. SA** | 1.64\|0.95 | 1.27\|0.58 | 1.71\|1.02 | **0.61**\|**0.17** |
| **Dis. LogP** | 0.37\|1.09 | 2.20\|2.93 | 1.44\|0.80 | **0.11**\|**0.63** |
| **Dis. Weight** | 181.4\|61.1 | 73.8\|**47.0** | 291.2\|170.5 | **46.3**\|74.4 |

**Table S5. The SMILES of Reference Active Compounds and 134 Selected Molecules Generated by AIxFuse**

| SMILES | Name |
|---|---|
| CC1=C(OC2CCN(C(CC(C(F)(F)F)C(F)(F)F)=O)CC2)C=C(Cl)C=C1NC3=NC4=C(CCC[C@H]4O)S3 | (R)-14d |
| Cc1ncc(C(=O)Nc2cc(Cl)cc(CN3CCN(C(=O)C4CCCC4)[C@@H](C)C3)c2C)cn1 | GSK-98E |
| COCCOc1cc(C#N)cc(C(=O)Nc2cnc3c(c(C4CCN(C(=O)C5CCCC5)CC4)cn3C)c2C)c1 | BDBM189939 |
| N#Cc1cccc(C(=O)Nc2cccc3c2CCC3N2CCN(C(=O)C3CCCC3)CC2)c1 | BDBM50509677 |
| CCn1c(CO)nn(-c2cc(O[C@@H](C)C(F)(F)F)c(cc2F)C(=O)Nc2c(F)cccc2Cl)c1=O | BAY2402234 |
| Cc1ccc(C(F)(F)F)cc1NC(=O)c1cc(F)c(-n2nc3n(c2=O)CCCC3)cc1OC(C)C(F)(F)F | BDBM490418 |
| Cc1ccc(F)cc1NC(=O)c1cc(F)c(-n2nc3n(c2=O)CCCC3)cc1OC(C)C1CCCCC1 | BDBM490490 |
| CCn1c(CO)nn(Cc2cccc(C(=O)Nc3cccc4c3CCC4N3CCN(C(=O)C4CCCC4)CC3)c2)c1=O | AF-1 |
| CC(C)Oc1cc(-n2nc3n(c2=O)CCCC3)c(F)cc1C(=O)Nc1cc(N2CCN(C(=O)C3CCCC3)CC2)ccc1C | AF-2 |
| CCOc1cc(-n2nc(CC)n(C)c2=O)c(F)cc1C(=O)Nc1cccc2c1CCC2N1CCN(C(=O)C2CCCC2)CC1 | AF-3 |
| CCn1c(CO)nn(-c2ccc3c(=O)n(-c4ccccc4CCC4CCN(C(=O)C(C)C)CC4)cc(C(C)=C)c3c2)c1=O | AF-4 |
| CC(Oc1cc(-n2nc3n(c2=O)CCCC3)c(F)cc1C(=O)Nc1cc(F)cc(N2CCN(C(=O)C3CCCC3)CC2)c1Cl)C(F)(F)F | AF-5 |
| CCCC(C)Oc1cc(-n2nc3n(c2=O)CCCC3)c(F)cc1C(=O)Nc1cc(F)cc(N2CCN(C(C)=O)CC2)c1Cl | AF-6 |
| Cc1c(NN=Cc2ccccc2C(O)=O)cnc2c1c(C1CCN(C(=O)C3CCOC3)CC1)cn2C | AF-7 |
| CCc1nn(-c2cc(OC(C)C)c(C(=O)Nc3ccccc3Cc3ccc(N4CCN(C(C)=O)CC4)cc3F)cc2F)c(=O)n1C | AF-8 |
| CCOc1cc(-n2nc3n(c2=O)CCCC3)c(F)cc1C(=O)Nc1cccc(F)c1CC1CCN(C(=O)C(C)C)CC1 | AF-9 |
| CCCCOc1cc(-n2nc(CC)n(C)c2=O)c(F)cc1C(=O)Nc1cccc(C(=O)N2CCN(C(=O)C3CCCC3)CC2)c1 | AF-10 |
| CCOc1cc(-n2nc3n(c2=O)CCCC3)c(F)cc1C(=O)Nc1cccc(C(=O)N2CCN(c3cccc(F)c3)CC2)c1C | AF-11 |
| CC(Oc1cc(-n2nc3n(c2=O)CCCC3)c(F)cc1C(=O)NC1(c2cc(Cl)cc(C3CC3)c2)CC1)C1CCCCC1 | AF-12 |
| CCc1nn(-c2cc(OC(C)C)c(C(=O)NC3(Cc4ccc(N5CCN(C(C)=O)CC5)cc4F)CC3)cc2F)c(=O)n1C | AF-13 |
| Cc1c(NC(=O)c2cc(F)c(-n3nc4n(c3=O)CCCC4)cc2O)cccc1C(=O)N1CCN(c2cccc(F)c2)CC1 | AF-14 |
| CCOc1cc(-n2nc3n(c2=O)CCCC3)c(F)cc1C(=O)Nc1cc(N2CCN(C(=O)C3CCCC3)CC2)ccc1C | AF-15 |
| CCOc1cc(-n2nc3n(c2=O)CCCC3)c(F)cc1C(=O)Nc1cccc(C(=O)N2CCN(C(=O)C3CCCC3)CC2)c1C | AF-16 |
| CCCCOc1cc(-n2nc3n(c2=O)CCCC3)c(F)cc1C(=O)NCCC1CCN(C(=O)C2CCCC2)C(C)C1 | AF-17 |
| CC(C)Oc1cc(-n2nc3n(c2=O)CCCC3)c(F)cc1C(=O)NC1CCN(C(=O)N2CCN(c3cccc(F)c3)CC2)CC1 | AF-18 |
| CC(C)Oc1cc(-n2nc3n(c2=O)CCCC3)c(F)cc1C(=O)Nc1ccccc1Cc1ccc(F)cc1 | AF-19 |
| CC(C)Oc1cc(-n2nc3n(c2=O)CCCC3)c(F)cc1C(=O)Nc1cccc(C2CCN(C(=O)C3CCCC3)C(C)C2)c1 | AF-20 |
| CC(=O)N1CCN(c2ccc(Cc3ccccc3NC(=O)c3ccc(-n4nc5n(c4=O)CCCC5)c(F)c3)c(F)c2)CC1 | AF-21 |
| CCCC(C)Oc1cc(-n2nc3n(c2=O)CCCC3)c(F)cc1C(=O)Nc1cccc(C(=O)C2CCCC2)c1Cl | AF-22 |
| CCOc1cc(-n2nc3n(c2=O)CCCC3)c(F)cc1C(=O)Nc1cccc(C(F)(F)c2ccccc2Cl)c1CO | AF-23 |
| CCOc1cc(-n2nc(CC)n(C)c2=O)c(F)cc1-c1ccc2[nH]cc(C3CCN(C(=O)C4CCCC4)CC3)c2c1 | AF-24 |
| CCc1nn(-c2cc(OC(C)C)c(C(=O)Nc3ccccc3CCc3ccccc3F)cc2F)c(=O)n1C | AF-25 |
| CCCCOc1cc(-n2nc(CC)n(C)c2=O)c(F)cc1C(=O)Nc1ccccc1Cc1cccc(F)c1 | AF-26 |
| CCCC(C)Oc1cc(-n2nc(CC)n(C)c2=O)c(F)cc1C(=O)NC1(c2ccc(N3CCN(C(C)=O)CC3)cc2F)CC1 | AF-27 |
| CCOc1cc(-n2nc3n(c2=O)CCCC3)c(F)cc1C(=O)C1CCN(C(=O)C2CCCC2)CC1C | AF-28 |
| CCOc1cc(-n2nc3n(c2=O)CCCC3)c(F)cc1C(=O)Nc1cccc(F)c1Cc1cccc(Cl)c1 | AF-29 |
| CCn1c(CO)nn(C(=O)Nc2cnc3c(c(C4CCN(C(=O)C(C)C)CC4)cn3C)c2OC)c1=O | AF-30 |
| CCc1nn(-c2cc(OC(C)C)c(C(=O)NC3(c4cc(Cl)cc(C5(CF)CC5)c4)CC3)cc2F)c(=O)n1C | AF-31 |
| O=C(Nc1ccccc1C(F)c1cccc(C(F)(F)F)c1)c1ccc(-n2nc3n(c2=O)CCCC3)c(F)c1 | AF-32 |
| CCCCOc1cc(-n2nc3n(c2=O)CCCC3)c(F)cc1C(=O)Nc1cccc(-c2ccc(F)cc2)c1C | AF-33 |
| CCc1nn(-c2cc(OC(C)C3CC3)c(C(=O)NC3(Cc4ccc(Cl)cc4C4CC4)CC3)cc2F)c(=O)n1C | AF-34 |
| CC(Oc1cc(-n2nc3n(c2=O)CCCC3)c(F)cc1C(=O)NC1(c2ccc(O)cc2F)CC1)C1CCCCC1 | AF-35 |
| CCc1nn(-c2cc(OC(C)C3CCCCC3)c(C(=O)NC3(c4cccc(C(F)(F)F)c4)CC3)cc2F)c(=O)n1C | AF-36 |
| CC(C)Oc1cc(-n2nc3n(c2=O)CCCC3)c(F)cc1C(=O)Nc1ccccc1CCc1cccc(F)c1 | AF-37 |
| CCOc1cc(-n2nc3n(c2=O)CCCC3)c(F)cc1C(=O)Nc1ccccc1Cc1cccc(C)c1F | AF-38 |
| CCc1nn(-c2cc(OC(C)C)c(C(=O)Nc3ccccc3C(F)(F)c3cccc(F)c3)cc2F)c(=O)n1C | AF-39 |
| CCc1nn(-c2cc(OC(C)C)c(C(=O)Nc3ccccc3CCc3c(F)cccc3F)cc2F)c(=O)n1C | AF-40 |
| CCCCOc1cc(-n2nc3n(c2=O)CCCC3)c(F)cc1C(=O)Nc1cc(C(F)(F)C(=O)C2CCCC2)ccc1C | AF-41 |
| CC(C)Oc1cc(-n2nc3n(c2=O)CCCC3)c(F)cc1C(=O)Nc1cccc(-c2cccc(F)c2)c1C#N | AF-42 |
| CCn1c(CO)nn(Nc2cnc3c(c(C4CCN(C(=O)CC(C)C)CC4)cn3C)c2C)c1=O | AF-43 |
| CCOc1cc(-n2nc3n(c2=O)CCCC3)c(F)cc1C(=O)Nc1ccccc1C(F)(F)c1ccc(F)cc1 | AF-44 |
| CCOc1cc(-n2nc3n(c2=O)CCCC3)c(F)cc1C(=O)Nc1cccc(C(F)(F)c2cccc(F)c2)c1CO | AF-45 |
| CCc1nn(-c2cc(OC(C)C)c(C(=O)Nc3ccccc3Cc3ccc(F)cc3)cc2F)c(=O)n1C | AF-46 |
| CCOc1cc(-n2nc3n(c2=O)CCCC3)c(F)cc1C(=O)Nc1ccc(F)cc1S(=O)(=O)c1cccc(Cl)c1 | AF-47 |
| O=C(Nc1cccc(C(F)(F)F)c1Cc1ccccc1F)c1ccc(-n2nc3n(c2=O)CCCC3)c(F)c1 | AF-48 |

| | |
|---|---|
| CC(C)(C(=O)Nc1cccc(F)c1)c1ccc(NC(=O)Cc2cc(F)c(-n3nc4n(c3=O)CCCC4)cc2O)cc1 | AF-49 |
| CCOc1cc(-n2nc3n(c2=O)CCCC3)c(F)cc1C(=O)Nc1cccc(C(F)(F)c2ccccc2F)c1CO | AF-50 |
| CCOc1cc(-n2nc3n(c2=O)CCCC3)c(F)cc1C(=O)Nc1cccc(-c2cccc(Cl)c2)c1Cl | AF-51 |
| CCOc1cc(-n2nc3n(c2=O)CCCC3)c(F)cc1C(=O)Nc1cccc(-c2cccc(Cl)c2)c1C | AF-52 |
| CC(C)Oc1cc(-n2nc3n(c2=O)CCCC3)c(F)cc1C(=O)Nc1cccc(-c2ccc(F)c(C)c2)c1CO | AF-53 |
| CCOc1cc(-n2nc3n(c2=O)CCCC3)c(F)cc1C(=O)Nc1cc(F)ccc1Cc1ccccc1F | AF-54 |
| CCOc1cc(-n2nc3n(c2=O)CCCC3)c(F)cc1C(=O)NC1CCC(F)(c2ccc(C3CC3)cc2Cl)CC1 | AF-55 |
| CCc1nn(-c2cc(OC(C)C)c(C(=O)NC3(c4cc(Cl)cc(C5(C(F)F)CC5)c4)CC3)cc2F)c(=O)n1C | AF-56 |
| CCn1c(CO)nn(Nc2cnc3c(c(C4CCN(C(=O)C(C)C)CC4)cn3C)c2C(F)(F)F)c1=O | AF-57 |
| CCOc1cc(-n2nc3n(c2=O)CCCC3)c(F)cc1C(=O)Nc1ccccc1Cc1ccc(F)cc1O | AF-58 |
| CCOc1cc(-n2nc(CC)n(C)c2=O)c(F)cc1C(=O)Nc1ccccc1Cc1ccc(F)cc1 | AF-59 |
| CCOc1cc(-n2nc3n(c2=O)CCCC3)c(F)cc1C(=O)Nc1cc(F)cc(-c2cccc(F)c2)c1Cl | AF-60 |
| CC(C)Oc1cc(-n2nc3n(c2=O)CCCC3)c(F)cc1C(=O)Nc1cccc(-c2ccccc2Cl)c1C#N | AF-61 |
| CCOc1cc(-n2nc(CC)n(C)c2=O)c(F)cc1C(=O)Nc1ccccc1C(F)(F)c1ccccc1F | AF-62 |
| CCc1nn(-c2cc(OC(C)c3ccccc3)c(C(=O)NC3(c4ccccc4Cl)CC3)cc2F)c(=O)n1C | AF-63 |
| CCOc1cc(-n2nc3n(c2=O)CCCC3)c(F)cc1C(=O)Nc1cccc(CCc2cccc(F)c2)c1C | AF-64 |
| CCc1nn(-c2ccc(C(=O)Nc3ccccc3CCc3cccc(Cl)c3)cc2F)c(=O)n1C | AF-65 |
| O=C(Nc1ccccc1C(F)(F)c1cccc(F)c1)c1ccc(-n2nc3n(c2=O)CCCC3)c(F)c1 | AF-66 |
| CCOc1cc(-n2nc(CC)n(C)c2=O)c(F)cc1C(=O)Nc1ccccc1CCc1cccc(CC)c1 | AF-67 |
| CC(C)Oc1cc(-n2nc3n(c2=O)CCCC3)c(F)cc1C(=O)Nc1cccc(-c2ccc(F)cc2)c1Cl | AF-68 |
| O=C(Nc1ccc(F)cc1Cc1ccc(F)cc1)c1ccc(-n2nc3n(c2=O)CCCC3)c(F)c1 | AF-69 |
| CCOc1cc(-n2nc3n(c2=O)CCCC3)c(F)cc1C(=O)Nc1cccc(-c2cccc(C(F)(F)F)c2)c1C#N | AF-70 |
| OC(=O)C1=C(C(=O)NCc2ccc(-c3ccc(C(O)(C(F)(F)F)C(F)(F)F)cc3)c(C3CC3)c2)CCC1 | AF-71 |
| CC(C)Oc1cc(-n2nc3n(c2=O)CCCC3)c(F)cc1C(=O)Nc1cccc(-c2ccc(F)cc2)c1C#N | AF-72 |
| Oc1cc(-n2nc3n(c2=O)CCCC3)c(F)cc1C(=O)Nc1ccccc1C(F)(F)c1cccc(Cl)c1 | AF-73 |
| Oc1cc(-n2nc3n(c2=O)CCCC3)c(F)cc1C(=O)Nc1cccc(C(F)(F)F)c1Cc1cccc(F)c1 | AF-74 |
| CCOc1cc(-n2nc3n(c2=O)CCCC3)c(F)cc1C(=O)Nc1cccc(-c2cccc(F)c2)c1C | AF-75 |
| COc1ccccc1C(=O)Nc1cnc2c(c(C3CCN(C(=O)C(C)C(F)(F)F)CC3)cn2C)c1C | AF-76 |
| OC(=O)C1=C(C(=O)Nc2ccc(-c3cccc(C(=O)C4CCCC4)c3)cc2F)CCC1 | AF-77 |
| CCc1nn(-c2cc(O)c(C(=O)Nc3ccccc3CCc3cccc(Cl)c3)cc2F)c(=O)n1C | AF-78 |
| CC(C)Oc1cc(-n2nc3n(c2=O)CCCC3)c(F)cc1C(=O)Nc1cccc(-c2cccc(Cl)c2)c1CO | AF-79 |
| CCc1nn(-c2ccc(C(=O)Nc3ccccc3Nc3cccc(C(F)(F)F)c3)cc2F)c(=O)n1C | AF-80 |
| CCOc1cc(-n2nc3n(c2=O)CCCC3)c(F)cc1C(=O)NC1CCC(F)(Cc2cccc(F)c2)CC1 | AF-81 |
| CCOc1cc(-n2nc3n(c2=O)CCCC3)c(F)cc1C(=O)Nc1cccc(-c2cccc(Cl)c2)c1CO | AF-82 |
| CCCC(C)Oc1cc(-n2nc(CC)n(C)c2=O)c(F)cc1C(=O)NC1(c2cccc(C(F)(F)F)c2)CC1 | AF-83 |
| CCOc1cc(-n2nc3n(c2=O)CCCC3)c(F)cc1C(=O)Nc1cccc(-c2ccc(F)cc2)c1C | AF-84 |
| Oc1cc(-n2nc3n(c2=O)CCCC3)c(F)cc1C(=O)Nc1ccccc1C(F)(F)c1ccc(F)cc1 | AF-85 |
| CCOc1cc(-n2nc(CC)n(C)c2=O)c(F)cc1C(=O)Nc1ccccc1C(F)(F)c1cccc(F)c1 | AF-86 |
| CCOc1cc(-n2nc3n(c2=O)CCCC3)c(F)cc1C(=O)Nc1cccc(C(F)(F)c2ccccc2F)c1 | AF-87 |
| CCOc1cc(-n2nc3n(c2=O)CCCC3)c(F)cc1C(=O)Nc1cc(F)cc(-c2ccccc2F)c1Cl | AF-88 |
| Cc1c(NC(=O)c2cc(F)c(-n3nc4n(c3=O)CCCC4)cc2O)cccc1C(F)(F)c1cccc(F)c1 | AF-89 |
| Oc1cc(-n2nc3n(c2=O)CCCC3)c(F)cc1C(=O)Nc1cccc(F)c1Cc1ccc(F)cc1 | AF-90 |
| Oc1cc(-n2nc3n(c2=O)CCCC3)c(F)cc1C(=O)Nc1cccc(F)c1Cc1cccc(F)c1 | AF-91 |
| CC(Oc1cc(-n2nc3n(c2=O)CCCC3)c(F)cc1C(=O)Nc1cccc(-c2ccc(F)cc2)c1CO)C(F)(F)F | AF-92 |
| Cc1c(NC(=O)c2ccc(-n3nc4n(c3=O)CCCC4)c(F)c2)cccc1C(F)(F)c1ccccc1Cl | AF-93 |
| CC(Oc1cc(-n2nc3n(c2=O)CCCC3)c(F)cc1C(=O)Nc1cccc(-c2ccccc2F)c1CO)C(F)(F)F | AF-94 |
| Oc1cc(-n2nc3n(c2=O)CCCC3)c(F)cc1C(=O)Nc1ccccc1C(F)Cc1ccccc1F | AF-95 |
| CCOc1cc(-n2nc3n(c2=O)CCCC3)c(F)cc1C(=O)NC1CCN(C(=O)c2ccc(C3CC3)cc2Cl)CC1 | AF-96 |
| Oc1cccc(-c2ccc(NC(=O)C3=C(C(=O)C4CCN(C(=O)C5CCCC5)CC4)CCC3)c(F)c2)c1 | AF-97 |
| CC(C)Oc1cc(-n2nc3n(c2=O)CCCC3)c(F)cc1CC(=O)Nc1ccc(C(C)(C)C(=O)Nc2ccc(F)cc2)cc1 | AF-98 |
| CCc1nn(-c2cc(O)c(C(=O)Nc3ccccc3C(F)(F)c3ccc(F)cc3)cc2F)c(=O)n1C | AF-99 |
| CCOc1cc(-n2nc3n(c2=O)CCCC3)c(F)cc1C(=O)Nc1cc(F)ccc1Oc1cccc(F)c1 | AF-100 |
| CCOc1cc(-n2nc3n(c2=O)CCCC3)c(F)cc1C(=O)NC1(c2ccc(-c3ccccc3F)cc2)CC1 | AF-101 |
| CCc1nn(-c2cc(O)c(C(=O)Nc3ccccc3Cc3cccc(C(F)(F)F)c3)cc2F)c(=O)n1C | AF-102 |
| Cc1c(NC(=O)c2cc(F)c(-n3nc4n(c3=O)CCCC4)cc2O)cccc1C(F)(F)c1ccccc1F | AF-103 |
| Oc1cc(-n2nc3n(c2=O)CCCC3)c(F)cc1C(=O)Nc1cccc(-c2cccc(F)c2)c1Cl | AF-104 |
| CCc1nn(-c2cc(OC(C)CC(=O)C3CCCC3)c(C(=O)Nc3ccccc3C(F)(F)F)cc2F)c(=O)n1C | AF-105 |
| CCOc1cc(-n2nc3n(c2=O)CCCC3)c(F)cc1C(=O)Nc1cccc(S(=O)(=O)Cc2ccc(F)cc2)c1CO | AF-106 |

| | |
|---|---|
| CCCCOc1cc(-n2nc3n(c2=O)CCCC3)c(F)cc1C(=O)Nc1ncccc1Cc1ccccc1F | AF-107 |
| O=C(NC1(c2ccc(-c3cccc(Cl)c3)cc2)CC1)c1ccc(-n2nc3n(c2=O)CCCC3)c(F)c1 | AF-108 |
| Oc1cc(-n2nc3n(c2=O)CCCC3)c(F)cc1C(=O)NC1(c2ccc(C3(CF)CC3)cc2Cl)CC1 | AF-109 |
| OCc1c(NC(=O)c2ccc(-n3nc4n(c3=O)CCCC4)c(F)c2)cccc1-c1cccc(C(F)(F)F)c1 | AF-110 |
| OCc1c(NC(=O)c2cc(F)c(-n3nc4n(c3=O)CCCC4)cc2O)cccc1C(F)(F)c1ccccc1Cl | AF-111 |
| CC(=O)CC(=O)Nc1ccc(-c2ccc(N(C)C(=O)c3c(F)cccc3Cl)c(OCC3CCC3)c2)cc1 | AF-112 |
| OCc1c(NC(=O)c2ccc(-n3nc4n(c3=O)CCCC4)c(F)c2)cccc1C(F)(F)c1ccccc1F | AF-113 |
| CC(C)Oc1cc(-n2nc3n(c2=O)CCCC3)c(F)cc1C(=O)NC1(c2ccc(-c3cccc(F)c3)cc2)CC1 | AF-114 |
| OCc1c(NC(=O)c2cc(F)c(-n3nc4n(c3=O)CCCC4)cc2O)cccc1-c1cccc(C(F)(F)F)c1 | AF-115 |
| OCc1c(NC(=O)c2cc(F)c(-n3nc4n(c3=O)CCCC4)cc2O)cccc1C(F)(F)C(=O)C1CCCC1 | AF-116 |
| CCOc1cc(-n2nc3n(c2=O)CCCC3)c(F)cc1C(=O)Nc1cccc(C(F)(F)C(=O)C2CCCC2)c1CO | AF-117 |
| CCOc1cc(-n2nc(CC)n(C)c2=O)c(F)cc1CC(=O)Nc1ccc(C(C)(C)C(=O)Nc2ccc(F)cc2)cc1 | AF-118 |
| CC(C)Oc1cc(-n2nc3n(c2=O)CCCC3)c(F)cc1C(=O)Nc1oncc1-c1cccc(C(F)(F)F)c1 | AF-119 |
| CC(C)(C(=O)Nc1ccccc1)c1ccc(NC(=O)Cc2cc(F)c(-n3nc4n(c3=O)CCCC4)cc2O)cc1 | AF-120 |
| OCc1c(NC(=O)c2ccc(-n3nc4n(c3=O)CCCC4)c(F)c2)cccc1C(F)(F)c1ccccc1Cl | AF-121 |
| Cc1cc(C(O)=O)c2[nH]c(-c3ccc(NCCc4ccc(C(F)(F)F)cc4)c(Cl)c3)nc2c1 | AF-122 |
| OC(c1ccc(-n2ncc3c2CCCC3NC(=O)c2noc3c2CCCC3)cc1)(C(F)(F)F)C(F)(F)F | AF-123 |
| Cc1c(NC(=O)c2cc(F)c(-n3nc4n(c3=O)CCCC4)cc2O)cccc1C(=O)c1ccccc1Cl | AF-124 |
| CCn1c(CO)nn(Nc2ccc(NCCc3ccc(C(F)(F)F)cc3)c(Cl)c2)c1=O | AF-125 |
| CCC(C)Oc1cc(-n2nc3n(c2=O)CCCC3)c(F)cc1C(=O)Nc1oncc1-c1cccc(C(F)(F)F)c1 | AF-126 |
| CC(Oc1cc(-n2nc3n(c2=O)CCCC3)c(F)cc1C(=O)NC1(c2c(Cl)cccc2C2CC2)CC1)C(F)(F)F | AF-127 |
| O=C(NC1CCCc2c1cnn2OCc1ccc(C(F)(F)F)cc1)c1noc2c1CCCC2 | AF-128 |
| CCn1c(CO)nn(Nc2ccc(C(C)(C)C(=O)Nc3cccc(F)c3)cc2)c1=O | AF-129 |
| OC(c1ccc(-n2ncc3c2CCCC3NC(=O)c2onc3c2CCCC3)cc1)(C(F)(F)F)C(F)(F)F | AF-130 |
| CCC(=O)CC(=O)Nc1ccc(-c2ccc(C(O)(C(F)(F)F)C(F)(F)F)cc2CC)cc1 | AF-131 |
| CC(C(C)=O)C(=O)Nc1ccc(S(=O)(=O)Cc2ccc(C3CC3)cc2Cl)cc1 | AF-132 |
| O=C(Nc1oncc1-c1cccc(C(F)(F)F)c1)c1ccc(-n2nc3n(c2=O)CCCC3)c(F)c1 | AF-133 |
| CCC(=O)CC(=O)Nc1ccc(-c2ccc(C(O)(C(F)(F)F)C(F)(F)F)cc2)cc1 | AF-134 |

Table S6. The MM/GSBA and ABFEP Binding Free Energies of Reference Active Compounds and 134 Selected Molecules Generated by AIxFuse

| Name | MM/GBSA △G(kcal/mol) | | | ABFEP △G(kcal/mol) | | | IC50(nM) | |
|---|---|---|---|---|---|---|---|---|
| | RORγt | DHODH | Sum | RORγt | DHODH | Sum | RORγt | DHODH |
| (R)-14d | -59.91 | -38.05 | -97.96 | -20.36 | -8.99 | -29.35 | 110 | 297 |
| GSK-98K(charged) | -62.66 | | | -6.54 | | | 0.7 | |
| GSK-98K(neutral) | -67.51 | | | -18.97 | | | 0.7 | |
| BDBM189939 | -74.79 | | | -21.63 | | | 12 | |
| BDBM50509677 | -70.27 | | | -19.92 | | | 465 | |
| BAY2402234 | | -50.40 | | | -17.09 | | | 4.17 |
| BDBM490418 | | -55.77 | | | -18.59 | | | 35 |
| BDBM490490 | | -64.72 | | | -18.82 | | | 6.85 |
| AF-5 | -74.44 | -66.55 | -140.99 | -18.68 | -21.45 | -40.31 | | |
| AF-20 | -60.57 | -61.76 | -122.33 | -17.48 | -18.02 | -35.50 | | |
| AF-2 | -75.46 | -71.27 | -146.73 | -16.01 | -17.93 | -33.94 | | |
| AF-16 | -72 | -54.34 | -126.34 | -17.65 | -15.82 | -33.47 | | |
| AF-3 | -76.97 | -65.76 | -142.73 | -13.92 | -18.07 | -31.99 | | |
| AF-1 | -83.33 | -63.52 | -146.85 | -20.00 | -6.19 | -26.19 | | |
| AF-4 | -70.95 | -71.45 | -142.4 | -6.88 | -14.05 | -20.93 | | |
| AF-8 | -70.54 | -67.32 | -137.86 | -15.22 | | | | |
| AF-10 | -70.2 | -66.1 | -136.3 | -14.23 | | | | |
| AF-13 | -68.68 | -64.27 | -132.95 | -15.22 | | | | |
| AF-15 | -69.12 | -57.3 | -126.42 | -13.11 | | | | |
| AF-17 | -62.04 | -63.81 | -125.85 | -16.33 | | | | |
| AF-21 | -65.64 | -55.99 | -121.63 | -12.81 | | | | |
| AF-24 | -59.37 | -60.08 | -119.45 | -15.69 | | | | |
| AF-27 | -58.11 | -60.19 | -118.3 | -8.58 | | | | |
| AF-28 | -58.2 | -59.7 | -117.9 | -9.83 | | | | |
| AF-49 | -55.46 | -54.51 | -109.97 | -13.43 | | | | |
| AF-6 | -71.68 | -67.78 | -139.46 | | | | | |
| AF-7 | -68.83 | -69.68 | -138.51 | | | | | |
| AF-9 | -72.5 | -65.02 | -137.52 | | | | | |
| AF-11 | -68.06 | -67.41 | -135.47 | | | | | |
| AF-12 | -60.06 | -74.65 | -134.71 | | | | | |
| AF-14 | -64.85 | -63.45 | -128.3 | | | | | |
| AF-18 | -61.8 | -63.12 | -124.92 | | | | | |
| AF-19 | -60.09 | -63.28 | -123.37 | | | | | |
| AF-22 | -56.85 | -64.27 | -121.12 | | | | | |
| AF-23 | -59.01 | -61.24 | -120.25 | | | | | |
| AF-25 | -54.62 | -64.81 | -119.43 | | | | | |
| AF-26 | -53.88 | -64.44 | -118.32 | | | | | |
| AF-29 | -57.68 | -59.45 | -117.13 | | | | | |
| AF-30 | -63.11 | -53.83 | -116.94 | | | | | |
| AF-31 | -55.67 | -61.07 | -116.74 | | | | | |
| AF-32 | -53.75 | -62.61 | -116.36 | | | | | |
| AF-33 | -48.69 | -66.96 | -115.65 | | | | | |
| AF-34 | -49.07 | -66.37 | -115.44 | | | | | |
| AF-35 | -55.22 | -59.87 | -115.09 | | | | | |
| AF-36 | -49.98 | -65.05 | -115.03 | | | | | |
| AF-37 | -54.67 | -60.19 | -114.86 | | | | | |
| AF-38 | -51.86 | -62.45 | -114.31 | | | | | |
| AF-39 | -49.51 | -64.79 | -114.3 | | | | | |
| AF-40 | -52.75 | -60.85 | -113.6 | | | | | |
| AF-41 | -53.36 | -60.03 | -113.39 | | | | | |
| AF-42 | -53.02 | -59.58 | -112.6 | | | | | |
| AF-43 | -55.3 | -57.07 | -112.37 | | | | | |
| AF-44 | -52.4 | -58.46 | -110.86 | | | | | |
| AF-45 | -53.59 | -57.2 | -110.79 | | | | | |
| AF-46 | -52.87 | -57.84 | -110.71 | | | | | |

| | | | |
|---|---|---|---|
| AF-47 | -45.07 | -65.6 | -110.67 |
| AF-48 | -52.31 | -57.78 | -110.09 |
| AF-50 | -49.2 | -60.37 | -109.57 |
| AF-51 | -51.36 | -58.17 | -109.53 |
| AF-52 | -53.58 | -55.83 | -109.41 |
| AF-53 | -50.9 | -58.46 | -109.36 |
| AF-54 | -50.88 | -58.09 | -108.97 |
| AF-55 | -56.43 | -52.15 | -108.58 |
| AF-56 | -51.25 | -57.21 | -108.46 |
| AF-57 | -52.05 | -56.18 | -108.23 |
| AF-58 | -54.14 | -53.94 | -108.08 |
| AF-59 | -51.16 | -56.85 | -108.01 |
| AF-60 | -51.26 | -56.57 | -107.83 |
| AF-61 | -51.78 | -55.88 | -107.66 |
| AF-62 | -47.2 | -60.45 | -107.65 |
| AF-63 | -47.3 | -60.3 | -107.6 |
| AF-64 | -44.69 | -62.73 | -107.42 |
| AF-65 | -45.67 | -61.69 | -107.36 |
| AF-66 | -46.35 | -60.6 | -106.95 |
| AF-67 | -51.06 | -55.8 | -106.86 |
| AF-68 | -50.38 | -56.46 | -106.84 |
| AF-69 | -47.81 | -58.42 | -106.23 |
| AF-70 | -51.36 | -54.71 | -106.07 |
| AF-71 | -60.97 | -44.96 | -105.93 |
| AF-72 | -51.82 | -53.68 | -105.5 |
| AF-73 | -50.38 | -55.11 | -105.49 |
| AF-74 | -50.04 | -55.41 | -105.45 |
| AF-75 | -49.78 | -55.33 | -105.11 |
| AF-76 | -58.09 | -46.76 | -104.85 |
| AF-77 | -48.35 | -56.31 | -104.66 |
| AF-78 | -52.36 | -52.3 | -104.66 |
| AF-79 | -49.01 | -55.63 | -104.64 |
| AF-80 | -51.69 | -52.73 | -104.42 |
| AF-81 | -47.37 | -57.03 | -104.4 |
| AF-82 | -45.16 | -58.92 | -104.08 |
| AF-83 | -47.73 | -55.54 | -103.27 |
| AF-84 | -49.11 | -54.16 | -103.27 |
| AF-85 | -48.54 | -54.4 | -102.94 |
| AF-86 | -47.26 | -55.1 | -102.36 |
| AF-87 | -46.76 | -55.12 | -101.88 |
| AF-88 | -45.94 | -55.52 | -101.46 |
| AF-89 | -44.65 | -56.6 | -101.25 |
| AF-90 | -47.92 | -53.21 | -101.13 |
| AF-91 | -48.81 | -52.3 | -101.11 |
| AF-92 | -46.7 | -54.16 | -100.86 |
| AF-93 | -50.58 | -50.17 | -100.75 |
| AF-94 | -46.31 | -54.39 | -100.7 |
| AF-95 | -47.59 | -53.09 | -100.68 |
| AF-96 | -48.58 | -51.21 | -99.79 |
| AF-97 | -51.68 | -47.87 | -99.55 |
| AF-98 | -49.83 | -49.67 | -99.5 |
| AF-99 | -44.74 | -54.76 | -99.5 |
| AF-100 | -45.43 | -54.05 | -99.48 |
| AF-101 | -43.09 | -56.22 | -99.31 |
| AF-102 | -49.27 | -49.12 | -98.39 |
| AF-103 | -44.65 | -53.35 | -98 |
| AF-104 | -44.51 | -53.43 | -97.94 |
| AF-105 | -54.87 | -42.76 | -97.63 |

| | | | |
|---|---|---|---|
| AF-106 | -53.65 | -43.85 | -97.5 |
| AF-107 | -42.82 | -54.66 | -97.48 |
| AF-108 | -44.92 | -52.3 | -97.22 |
| AF-109 | -49.07 | -48.14 | -97.21 |
| AF-110 | -47.45 | -49.51 | -96.96 |
| AF-111 | -44.42 | -52.39 | -96.81 |
| AF-112 | -52.64 | -43.97 | -96.61 |
| AF-113 | -43.68 | -52.9 | -96.58 |
| AF-114 | -49.85 | -46.62 | -96.47 |
| AF-115 | -42.23 | -54.04 | -96.27 |
| AF-116 | -40.61 | -55.28 | -95.89 |
| AF-117 | -38.17 | -55.7 | -93.87 |
| AF-118 | -54.73 | -38.95 | -93.68 |
| AF-119 | -43.64 | -49.14 | -92.78 |
| AF-120 | -45.31 | -47.01 | -92.32 |
| AF-121 | -42.44 | -49.6 | -92.04 |
| AF-122 | -48.14 | -43.34 | -91.48 |
| AF-123 | -48 | -42.22 | -90.22 |
| AF-124 | -39.98 | -49.7 | -89.68 |
| AF-125 | -42.76 | -46.61 | -89.37 |
| AF-126 | -35.73 | -53.01 | -88.74 |
| AF-127 | -34.65 | -53.75 | -88.4 |
| AF-128 | -45.52 | -42.73 | -88.25 |
| AF-129 | -39.27 | -48.97 | -88.24 |
| AF-130 | -46.64 | -31.69 | -78.33 |
| AF-131 | -36.75 | -41.31 | -78.06 |
| AF-132 | -37.38 | -38.38 | -75.76 |
| AF-133 | -36.41 | -38.62 | -75.03 |
| AF-134 | -37.17 | -36.42 | -73.59 |
